# Codon usage bias maps ocean microbiome adaptation to temperature-nutrients regimes

**DOI:** 10.64898/2026.10.02.755941

**Authors:** Corentin Gouzien, Antoine Bagnaro, Laurence Garczarek, Mathieu Ardyna, Fabien Joux, Samuel Chaffron

## Abstract

Microbial communities underpin ocean productivity and biogeochemistry, but their functional adaptation to environmental gradients remains poorly understood. Codon usage bias (CUB), a marker of translational efficiency, constrains growth rate potential, yet its role in functional adaptation across natural communities is underexplored. By analysing 589 metagenome-derived microbial communities from the global ocean, we demonstrate that CUB in 320 metabolism-related gene groups capture genomic adaptation to distinct temperature and nutrient concentrations. We identified four adaptation signatures shaped by distinct temperature-nutrient regimes, revealing how microbes optimize protein synthesis in their ecological niche. This genome-centric approach deepens our understanding of functional adaptation in uncultured marine microbes and provides a framework to inform ocean biogeochemical models and predict microbial responses to climate-driven changes.

## Introduction

Marine microbes drive half of global primary production (*1*), but predicting their functional adaptation and responses to environmental change remains a challenge. Common approaches to characterize ecological roles and strategies within natural microbial communities (*2*–*4*) leverage metabarcoding for species composition and reference genome annotations for prokaryotic traits inference (*5*). Nevertheless, so far very few studies have leveraged shotgun metagenomics for direct inference of functional traits (*6*). While functional traits are usually inferred at genome level, they can also be measured at various levels of organization, from species to communities. Such community-aggregated traits can be particularly useful to transform the ever-increasing amount of metagenomic data into ecological knowledge (*7*), which can subsequently inform mechanistic ecosystem models with microbial trait features (*8*).

Among prokaryotic phenotypic traits usually studied, such as motility, oxygen or pH tolerance, growth rate is one of the most available as it is relatively easy to measure. At the cellular level, growth rate corresponds to the number of cell divisions per unit of time. At the molecular scale, growth rate is the outcome of a symphony of many complex and interconnected processes (e.g., chromosome replication, ribosome synthesis, transcriptional regulation, protein synthesis). Selection for fast growth leaves traces of adaptation that significantly imprints prokaryotic genomes (*10*). One of such traces is ribosomal Codon Usage Bias (rCUB) (*11, 12*), which is based on codon frequencies in genes encoding ribosomal proteins. Codons used at frequencies matching expected frequencies will result in a low rCUB, while usage of few candidate codons diverging from expected, will result in a high rCUB. CUB can usually be explained by transfer ribonucleic acid (tRNA) availability in the cell (*10*). Codon decoded by highly available tRNAs are overrepresented in biased genes, hence accelerating protein elongation. Thus, for a given amino acid, distinct codons can lead to different speed in protein synthesis. This bias can also be related to messenger RNA (mRNA) stability (*13*–*15*). Ribosomes slow down when decoding non-optimal codons, inducing ribosome staling or recruiting DEAD-box helicase proteins, two mechanisms inducing mRNA decay (*15, 16*). mRNA stability may control the success of protein synthetized given a pool of mRNA.

Overall, by influencing the speed and reliability of protein synthesis, CUB is a marker of protein quantity in the cell, as shown in earlier studies (*11*). Although the interdependent molecular mechanisms underlying this pattern are still being investigated and debated (*17*), there is a clear consensus on the capacity of CUB to influence protein synthesis. We thus hypothesized that CUB can be a key marker of physiological and functional adaptation to environmental conditions (*18, 19*). To test this hypothesis, we first investigated the distribution of ribosomal Community Bias (rCoB, i.e., rCUB averaged at the community level) in the global ocean. Considering the link between rCUB and growth rate, rCoB provides a community-level proxy of potential growth that can be compare with other growth rate proxies (*3*). We assembled a large collection of Metagenome-Assembled Genomes (MAGs) profiled from 589 metagenomes at the surface of 334 sites covering all marine biomes in the global ocean. After mapping rCoB in the global ocean, we used statistical modelling to identify the main environmental drivers of potential growth (temperature vs. nutrient concentrations). Next, we went beyond ribosomal genes rCUB by investigating CUB in other functional groups of genes (fCUB). To our knowledge, this is the first large-scale exploration of functional CUB in natural microbial communities. We show that functional Community Bias (fCoB, i.e., fCUB averaged at the community level) captures signatures of microbial adaptation to different environmental conditions. Finally, we move from communities back to genomes and show that fCUB can be a key tool to characterize intra-genus variability and functional adaptation to distinct ecological niches.

## Results

### Environmental drivers of ocean microbiome rCoB

We leveraged global ocean metagenomes from three large oceanographic campaigns: *Tara* Oceans, Bio-GO-SHIP and GeoTraces (*20*–*22*). We focused our analyses on the euphotic layer (0-200m deep), and kept 589 metagenomes characterised by qualitative measurements of temperature and nutrient concentrations (n= 105, 154 and 330 per campaign, respectively). Using the MGnify marine genome catalogue (v2) with 13k Metagenome-Assembled Genomes (MAGs) dereplicated at species level (95% Average Nucleotide Identity), we profiled MAGs abundance and identified 737 unique MAGs on average per metagenome (with a standard deviation of 382). It revealed a good community representativeness but a relatively high heterogeneity between samples.

We then computed rCUB for all MAGs by comparing the codon frequencies in all ribosomal genes to expected codon frequencies (see Materials and Methods). Using the MAG abundance in each metagenome, we then computed rCoB, the community average of ribosomal bias (Figure 1-A and Materials and Methods). By giving higher weight to more abundant MAGs, we limit methodological bias (i.e., lower sequencing depth is mitigated) between campaigns (see Materials and Methods).

**Figure 1:**
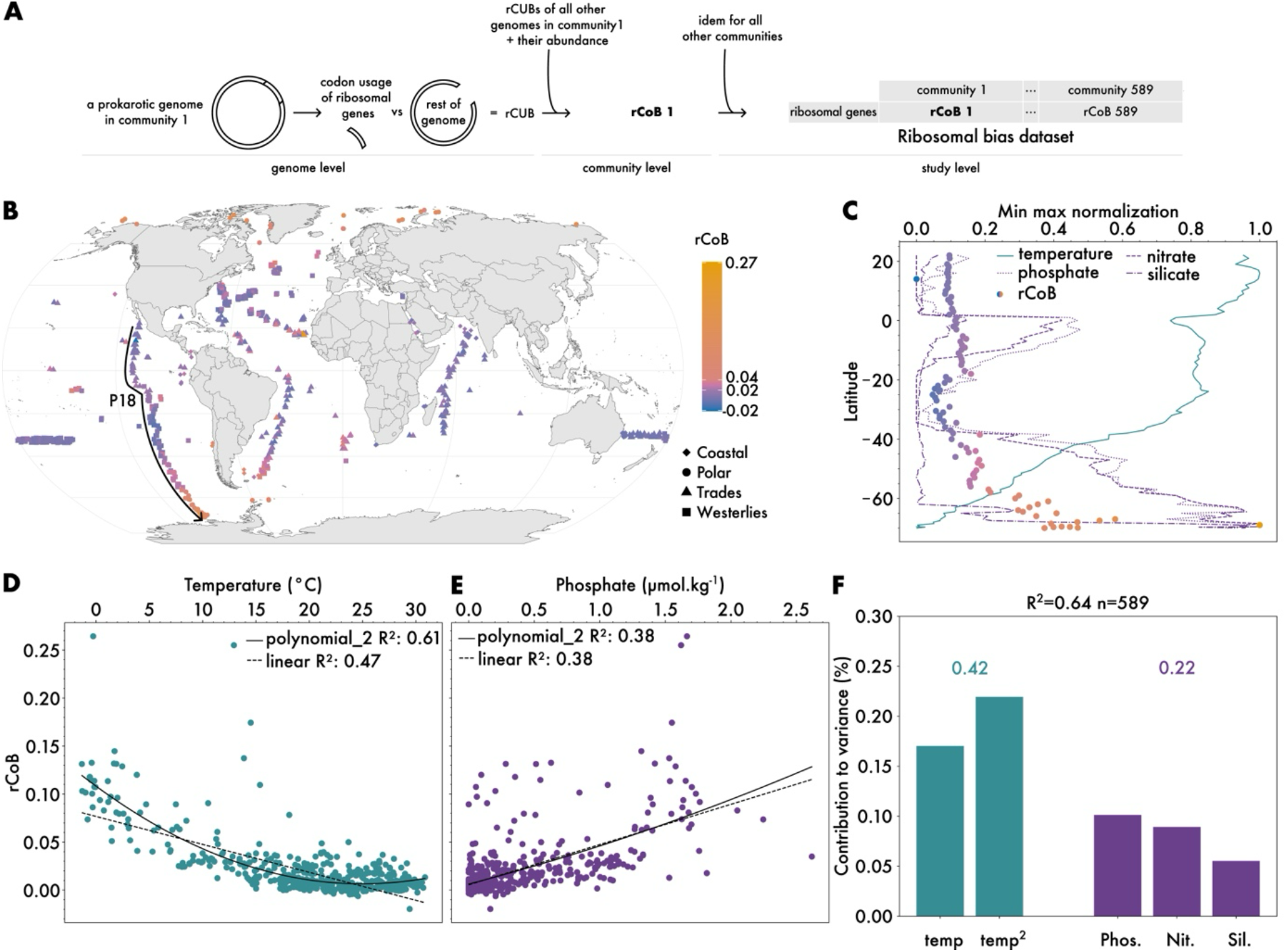
Environmental drivers of rCoB distribution, a proxy of community growth rate. **A**. Overall approach to compute rCoB (n=589) from genome-level CUB. **B**. Map of rCoB across the global ocean. Note that a small jitter was added to longitude and latitude in order to distinguish communities with similar coordinates but different depths. **C**. Temperature and nutrients are anti-correlated at the global and local scale of the transect P18 (labelled on B). Min max normalisation allows to scale different variables on the same plot, by affecting 1 to the maximum of the variable, and 0 to the minimum. Note a local drop of temperature around latitude 0, concomitant with a local peak of nutrients. There is a global tendency of nutrient to rise toward the southern pole with a tendency of temperatures to drop. rCoB are coloured based on their values in B. **D**. Correlation between temperature and rCoB (polynomial fit R^2^=0.61, p-value<10^−15^; linear fit R^2^= 0.47, p-value<10^−15^). **E**. Correlation between phosphate concentrations and rCoB (polynomial fit R^2^=0.38, p-value<10^−15^; linear fit R^2^= 0.38, p-value<10^−15^). Nitrate and silicate harbour similar correlations with rCoB. **F**. Variance explained by temperature and nutrient concentrations for the distribution of rCoB in a multiple linear regression model. Since we considered a relationship between temperature and rCoB as polynomial of order two, temperature has two components in the linear model, temp and temp2, the linear and quadratic component of the polynomial fit (see methods for more details). Pho., Nit. and Sil. = Phosphate, Nitrate and Silicate concentrations.

Mapping the rCoB at the global scale revealed a clear association with absolute latitude (Figure 1-BC). Overall, the pattern follows expected temperature and nutrient gradients (*23*), where we find cold, nutrient-rich waters at the poles compared to warm, nutrient-poor waters at the equator. Locally, high rCoB appeared to be associated with relative high nutrient concentrations and low temperatures. This pattern is particularly visible along the Bio-GO-SHIP P18 cruise that crossed the South Pacific Gyre, where we observe alternating rCoB following local variations in temperature and nutrients concentrations (Figure 1-BC). This pattern is also visible next to Cape Verde Island at a place where a relatively high rCoB occurs simultaneously with relatively high nutrients and low temperatures (Figure S1). Hence, rCoB is significantly shaped by gradients of temperature and nutrients concentrations.

The well-known and observed collinearity between temperature and nutrients concentrations at global and local scale, complicates the causal understanding of factors driving the distribution of rCoB. Do we find high rCoB in the poles because of local low temperatures or local high nutrient availability? To answer this question, we evaluated the links between rCoB, temperature and nutrients concentrations. By fitting a second order polynomial relationship, we improved the linear fit between rCoB and temperature (Figure 1-D). This was not the case for nutrients concentrations (Figure 1-E). Temperature displayed a negative correlation with rCoB, suggesting a negative relationship between potential community growth rate and temperature. It confirms previously reported relationships when considering Mean Copy Number (MCN) of the ribosomal RNA gene operon, another proxy of growth rate (*3*). However, the rCoB metric is likely more precise and robust than MCN as it is not based on metabarcoding data (involving potentially biased PCR amplification) and less sensitive to few individuals, even when adjusting MCN by improving its trait taxonomic assignation (Supplementary Text S1 and Figure S2, S3).

We next accounted for potential confounding effects, which is common between temperature and nutrient concentrations. Hence, these variables were integrated into a multiple linear regression model explaining the distribution of rCoB (Supplementary Text S2). We applied the model to five datasets: a global ocean model including all 589 metagenomes and four subsets, one for each of the four Longhurst marine biome defined by physical forcing (*24*) (Supplementary Text S3, Figure S4). For the global ocean model, nutrients and temperature explained 64% of the total variability of rCoB. The Lindeman, Merenda, and Gold (LMG) (*25*) contribution analysis further indicated that two third of this total variability is explained by temperature, highlighting its role as the main driver of rCoB in the global ocean (Figure 1-F). Nevertheless, nutrient concentrations explained a third of the total variance of the model, highlighting their significant and non-negligible impact on rCoB at global scale. At marine biome scale, nutrients effects can even predominate where temperature tends to be stable across seasons (*26*) (Supplementary Text S3, Figure S4).

Focusing on rCoB allows us to study the genotype component of the genotype–environment interaction driving an observed phenotype (*27*). Here we only study a potential maximum growth rate, which is likely different from the effective growth rate constrained by environmental factors. It was reported that the effective bacterial growth in the Arctic Ocean is three time lower than in the Equatorial Pacific (*28*). In contrast, potential growth rate proxied by rCoB suggests the opposite (Figure S5). This likely reflects a genomic adaptation to compensate for slower growth at lower temperature as discussed previously (*10, 29*). By studying ocean microbiome rCoB at global scale, we show a strong effect of not only temperature but also nutrients concentration in shaping genomic adaptation related to microbial growth strategies in the ocean. This is confirming previous results relating growth strategies to temperature using metabarcoding data only (*3*), but goes beyond by demonstrating a significant effect of nutrients on potential growth, while allowing to identify environmental genomes and functions driving the ecology and biogeography of marine microbes.

### Microbiome codon usage bias reveals functional signatures of adaptation

Given the strong genomic signature related to growth strategies (rCUB) observed in marine microbial genomes, we thought of exploring the CUB of other genes and associated functions (fCUB). To do so, we generalised the approach presented above by examining bias across 320 metabolism-related gene groups (i.e., KEGG modules labelled M00xxx (30), Figure 2-A). For each function, we compared the median bias of the group of gene of interest to the rest of the genome (Figure 2-A, Material and Methods).

**Figure 2:**
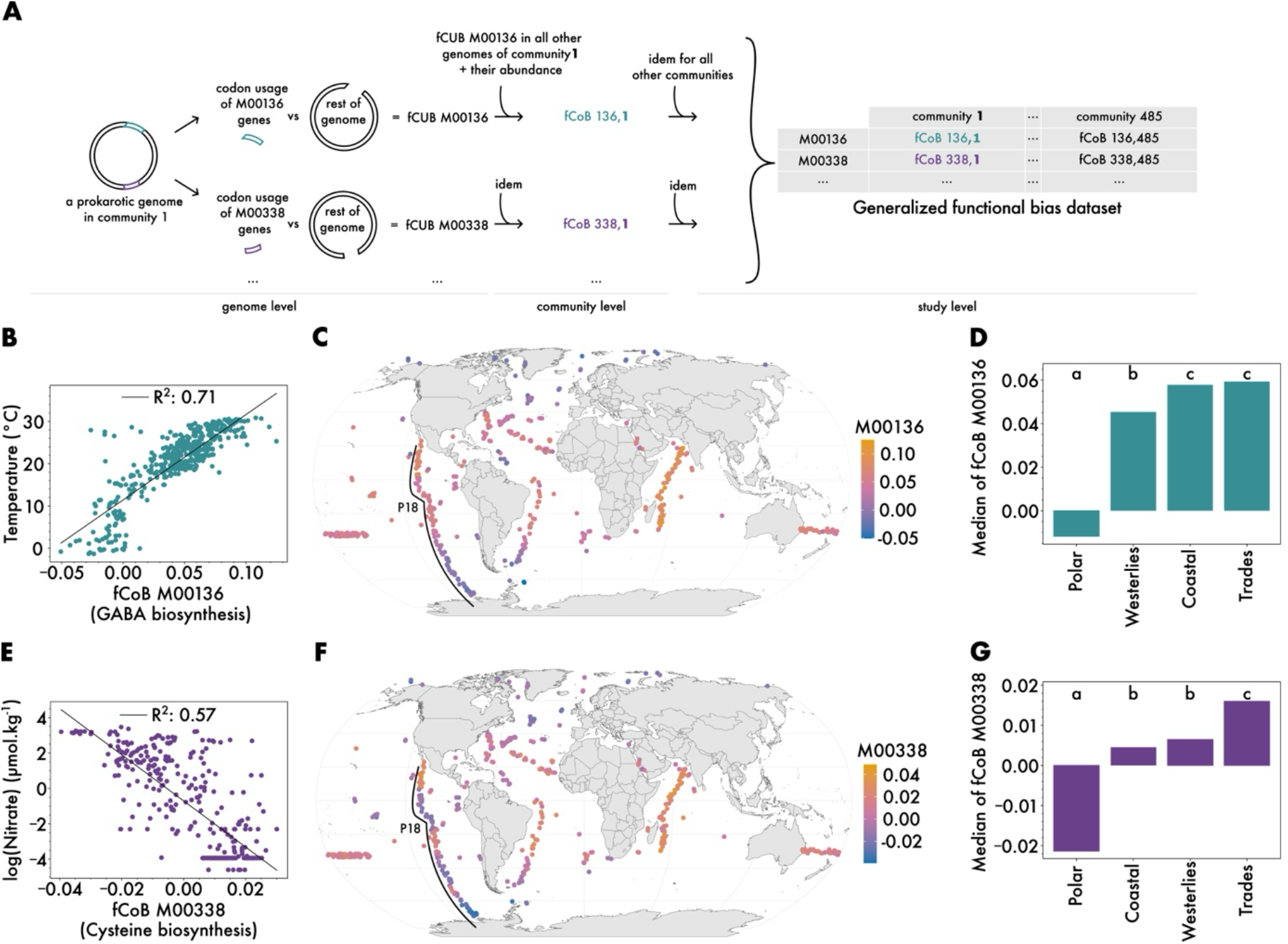
Functional Community Biases (fCoBs) reveal distinct signatures of genomic adaptation to environmental gradients at global scale. **A**. Computational framework to generalise CUB to functional community bias (fCoB). From previously one group of interest (ribosomal genes, Figure 1), we expand to 320 quasi-universal functions to characterize ocean microbial communities. **B**. Correlation between fCoB of KEGG module (*30*) M00136 and temperature **C**. Map of community bias of M00136 across the global ocean. Note that a small jitter was added to distinguish communities with similar coordinates but different depths. **D**. Median of community bias can serve as biome biomarkers. Letter represent statistically different medians (permutation test, p-value<0.05, n=10k) **E.F.G**. Idem as B.C.D. panel but considering the correlation between community bias of M00338 and nitrate concentrations. Note the differences between patterns along the P18 transect on C. and F. implying that we capture different signals even if temperature and nutrient are anti-correlated (Figure 1-C).

These fCUB can be averaged at the community level, which we named functional Community Bias (fCoB), similar to rCUB averaged into rCoB. The overall rational is that CUB can be a marker of functional adaptation in genomes. If a function is important for a given microbial genome and its community due to environmental pressures, then the whole group of genes related to this function will likely be biased too. Synchronized codon usage between functional related genes validate the pertinence of studying CUB per functional group (*17, 31*). Over the 320 considered functional modules, 39% were universal, i.e., were detected in all metagenomes of this study. The remaining 61% were quasi-universal, being detected on average in 96% of communities.

We explored the associations of the 320 functional biases with key environmental parameters. Some fCoB showed no correlation, such as module M00035, indicating no CUB significant variations in methionine degradation pathways across communities (Figure S6). On the contrary, the bias of seven modules showed strong and significant correlation with temperature or nitrate concentration, revealing highly environment-dependent fCoB (Figure 2, Figure S7). Notably, the biases of two specific modules are strongly associated with known environmental gradients, both particularly visible along the P18 transect (Figure 2-BCDEFG):

(i) The bias of M00338 captures nitrate-rich and nitrate-poor zones, being more biased in low-nitrate regions of the globe. Interestingly, M00338 is a functional module involved in cysteine synthesis during which ammonia is released in the cell (*30*). The genomic optimisation of this module may thus be a way to boost intra-cell nitrogen fluxes in nitrogen-poor waters.

(ii) The bias of M00136 captures the latitudinal temperature gradient. We suggest that the tight coupling between temperature and M00136 fCoB could be explained by a strict control over intracellular polyamine concentration. In prokaryotes, growth is inhibited by both excess and depletion of polyamine. Hence, its concentration in cells is tightly controlled (*32*). Warm waters could increase intracellular polyamines concentrations, either directly or indirectly, because of higher effective growth rates occurring in warm oceanic regions (*28, 33, 34*). Converting excess polyamine into gamma-aminobutyric acid (GABA), the pathway under the M00136 label, would be a way to detoxify cells in warm waters. A very high bias may be detrimental in region with lower effective growth rate, with too high levels of polyamines entering the GABA pathway, leaving too few molecules for basic metabolism. On the contrary, a low bias would limit the detoxification and would thus be detrimental in region with high effective growth rate given the toxicity of polyamines excess. Thus, a precise equilibrium of growth-produced polyamines may explain the strong correlation between temperature and fCoB of M00136.

Among the five other high correlating functions with environmental variable (Figure S7), we identified functions related to the pentose bisphosphate pathway, to threonine, glutathione and two lysine biosynthesis pathways. Lysine is a key component of proteins but also of peptidoglycans forming bacterial the cell wall (*35*), and is quite expensive to synthesize as compared to other cell wall integrated amino acids (*36, 37*). This may explain why we find high biased lysine synthesis communities in low-nitrate waters, given the central role of this amino acid for bacterial growth.

We described and discussed the fCoB in two metabolic modules as they showed very strong correlations with environmental parameters at global scale. This illustrates well the potential to explore the bias of any given function (e.g., amino acids and vitamins biosynthesis, iron-related metabolism) to better understand its environmental drivers in a given environment (e.g., low light, low oxygen concentration). Moreover, we show that the community bias (fCoB) in specific functions captures distinct patterns. Thus, by selecting a given set of functions with informative biases (i.e., biased signature functions) we hereinafter propose a new way to characterise the functional ecology of natural microbial communities using solely genomic information.

### Functional CUB reveals adaptative signatures to distinct temperature-nutrient regimes

Given the strong associations between fCoB and key environmental parameters reported above, we next evaluated the relevance of fCoBs to characterize the genomic adaptation and functional ecology of marine microbial communities. Indeed, functional biases aggregated at the community level could be particularly useful to capture both global and local signatures of functional adaptation across the global ocean. To this end, we projected 485 euphotic communities using surface metagenomes (0 to 100m deep), described by their biases in 320 quasi-universal functions (i.e., 320 fCoB), into a Uniform Manifold Approximation and Projection (UMAP) (Figure 3-AB, Figure S8). Here, this dimensionality reduction approach will cause communities with similar bias for a given function to be close to each other in the UMAP, whereas communities with opposite bias in a given function will be further apart in the projection.

**Figure 3:**
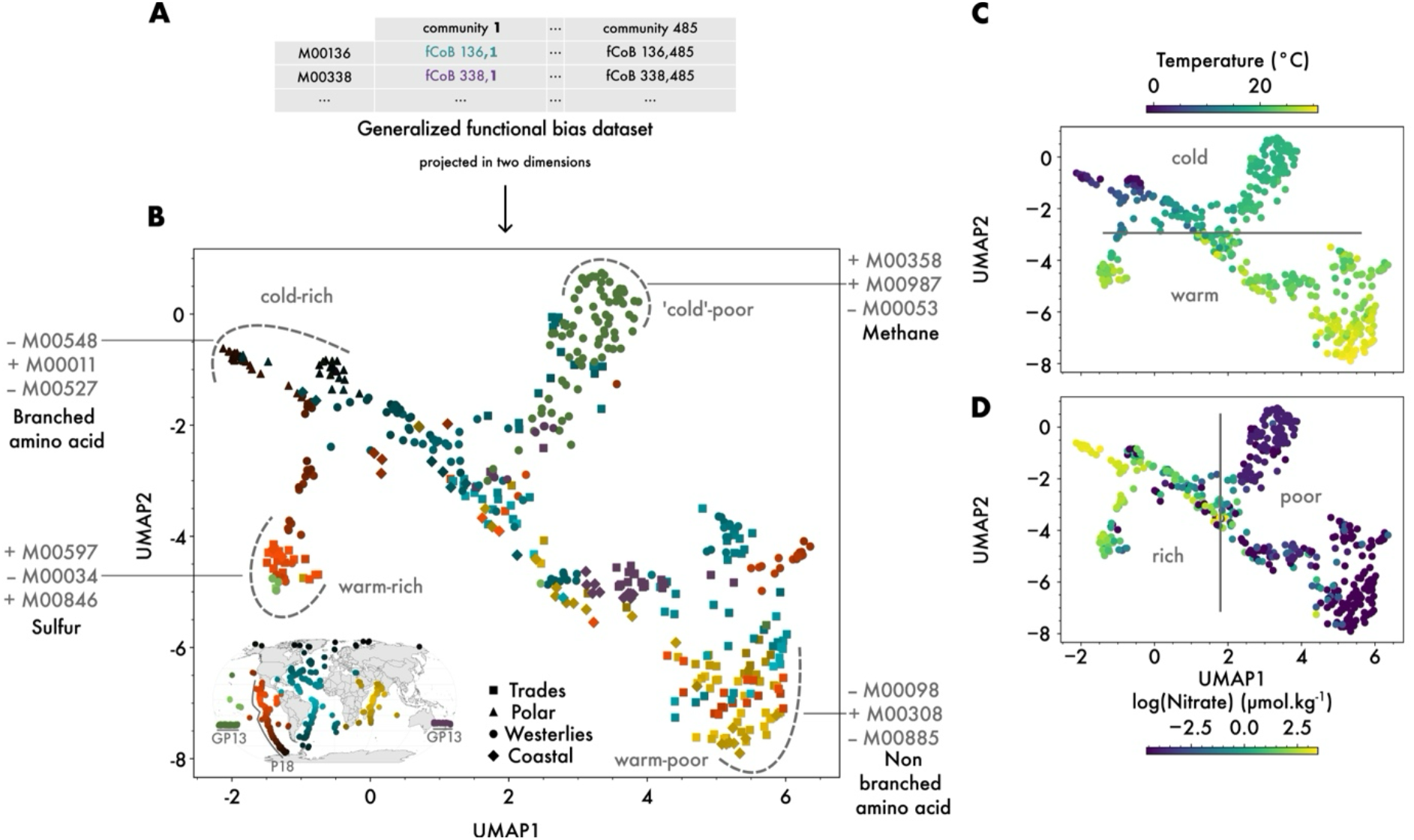
fCoBs reveal adaptative signatures to four distinct temperature-nutrient regimes. **A**. Dataset used in this plot is the same as in figure 2A. **B**. Uniform Manifold Approximation and Projection (UMAP) representing 485 communities according to their fCoBs. Colours help to visualize the geographical origin of communities by regrouping them by similar longitudes (colour gradient) and by similar brightness according to absolute latitude. Hence, darker colours represent polar communities, brighter colours equatorial communities, blue gradients Atlantic Ocean communities, and yellow gradients Indian Ocean communities. Note that colours are mixed across the UMAP, meaning that communities do not cluster solely based on their geographical origin. We visually identified clusters based on the shape of the plot, and labelled them (e.g., cold-rich, warm-poor) according to the tendency of temperature and nutrients concentration values from plot C.D. Note that in the ‘cold’-poor cluster, temperature is not that cold as compared to polar communities. For each cluster, we identified and highlighted three representative KEGG modules (e.g., M00009). + and – indicate if the module has a negative or positive bias in this cluster. **C**. Each community is coloured by its temperature showing how this variable explains cluster distribution. **D**. Idem as C. for nutrients. Only nitrate is shown but phosphate and silicate present similar distributions (Figure S9). Natural log scale was used, and when nutrient concentration was null, we imposed the colour to be equal to the log of the lowest non null value.

In this analysis, communities are only described by their functional bias values and no environmental data are used to obtain the UMAP projection. Nevertheless, temperature and nutrients concentrations largely explained the samples distribution in the UMAP (Figure 3-BCD, Figure S10), forming four clear clusters grouping communities belonging to four distinct temperature-nutrients regimes. Thus, this reveals conserved functional signatures of genomic adaptation selected by environmental constraints on marine prokaryotic communities. Notably, both polar communities (Arctic and Antarctic) cluster together, highlighting they have similar functional biases selected by polar environmental conditions (Figure 3-B). It also confirms their previously known similarity in terms of structure and function for taxa or genes (*38, 39*). Nevertheless, here similarity is not based on usual biomarkers but rather on similar optimisation patterns of functional gene groups. In other words, we explore orthologous genes variability with a relatively simple method (i.e., counting codons), and are able to show how this variability is a good marker for ecological niche adaptation. The scattering of the P18 transect across different clusters also strengthens the idea that communities cluster per temperature-nutrient regimes rather than by basin or oceanic region. Nevertheless, apart from this general tendency, some communities did not fit within a given temperature-nutrient regime, which is likely due to other factors driving bias selection but currently not available in our dataset (latent variables). For instance, it seems that depth, or co-varying factors sur as light availability, could be selecting for different functional bias signatures (Figure S11).

Next, to complement global functional signatures discussed previously (Figure 2), we explored cluster-specific local signatures by investigating functions uniquely biased in a given cluster (Figure 3-B). These functional modules present either positive or negative significative bias compared to other clusters. As CUB has been linked to protein dosage (*11, 17*), we can expect selection toward low or high dosage in specific clusters. On the one hand, having a high positive bias may be a way to enhance translational efficiency of proteins and thus eventual higher fluxes in a given metabolic pathway. On the other hand, having a negative bias may be a way to reduce translation rates and protein production, thereby reducing fluxes of a given metabolic pathway. To identify such signatures of bias, we ran an extreme gradient boosting (XGBoost) algorithm to learn which functions, based on their fCUB, were most useful to discriminate clusters (*40*). Here we focused on local cluster specific functions and thus did not consider functions showing global correlations (cf. Figure 2). Selected functions were ranked by total gain, quantifying the importance of the function in separating communities into distinct clusters. And representative functions were retained when their cluster-specific bias was visually supported (Figure 3-B, Figure S12). We did not intend to summarize the diversity of metabolic adaptation by focusing only on few representatives, but rather to reveal or confirm insights about these potential adaptations.

Considering the four different clusters identified, the *cold-rich* cluster comprising the coldest communities in the study, showed low bias for the synthesis of all amino acids (e.g., M00527 for lysine synthesis) excepted for branched-chain amino-acids (BCAAs). Intriguingly, BCAAs are precursors to branched-chain fatty acids known to increase membrane fluidity and allow growth in cold environments (*41*). Supporting this idea, we found positively biased oxaloacetate (M00011), a BCAA precursor, produced during the TCA cycle (i.e., Krebs cycle). High bias in BCAA and their precursors compared to low bias in other amino acids might be a way to direct amino acids building material toward mandatory cold related amino acids.

The *‘cold’-poor* cluster harboured high biases in coenzyme M (M00358) a methanogenesis related molecule, and sulfur-related metabolism (M00987). Given the need for a sulfur atom in coenzyme M, we suggest that this sulfur-related biased function may serve indirectly methane metabolism (*42*). The basis for the methanogenesis-associated signal in this cluster remains unclear.

Several mechanisms have been proposed for methane production in the open ocean, including methylphosphonate (Mnp) degradation or the anoxic zooplankton gut environment to methane production by prokaryotes (*43*). As most of the communities from this cluster come from the eastern part of the GP13 transect (Figure 3-A), and given that this region is not particularly methane-producing (*43*), we hypothesize that this methane-related specificity may correspond to a lower-rate metabolic pathway that is nonetheless essential for the development of prokaryotic communities in these relatively cold and nutrient-poor regions.

The *warm-poor* cluster contains low biases in acylglycerol degradation. In our analyses, it is actually the function with the highest *total gain* of all selected function, meaning this low bias is very typical of warm-poor regions. When investigating this degradation pathway, we found it relies here on the K01046 enzyme, a triacylglycerol (TAG) lipase. Limiting TAG degradation, constituent of lipid droplets, allow survival in harsh conditions (*44*). The importance of TAG in these communities is also supported by limiting other lipid synthesis like the *warm-poor* typical low biased mycolic acid synthesis (M00885). In addition, we find high bias in non BCAAs, opposed to the cold-rich cluster, probably to limit membrane fluidity in warm regions.

The *warm-rich* cluster is characterized by sulfuric function, potentially linked to the upwelling or island mass effect peculiar to this cluster (*45, 46*). Indeed, we find positively biased anoxygenic respiration based on sulfur (*47*), positively siroheme biosynthesis, which allows sulfur assimilation (*48*), and negatively biased methionine salvage pathway that recycles sulfur (*49*).

Delving into cluster-specific functions shed light on adaptation signatures for the corresponding microbial communities. While, *warm-rich* and *‘cold’-poor* communities showed respectively sulfur and methane metabolism biases, the warm-poor and cold-rich communities point towards biases in BCAA synthesis metabolism. These cluster bias specificities actually precise the global pattern of fCoBs (Figure 2). Overall, the generalised fCoB approach can reveal which pathways have likely been selected for adaptation to the environment and could also inform us on the dosage of cellular machinery in natural microbial communities. While wet lab experiments will be needed to better understand the consequences of biased or unbiased functions, few hypothesizes could explain fCoB variations. First, they may contribute to metabolite equilibrium in cells as suggested for polyamines. (Un)biased functions may also contribute to hierarchize or prioritize pathways sharing substrates or competing for energy, as seen with amino acid synthesis in cold-rich environments where BCAA appears to be prioritized over other amino acids. Such biases may also be useful to quickly adapt to sudden events, such as environmental stresses or nutrient availability, thanks to the rapid translation of key enzymes. Nevertheless, this CUB-based approach constitutes a simple and powerful method to characterize microbiome communities by identifying key functional adaptation to environmental conditions solely on the basis of (meta)genomics information.

### Functional CUB identifies signatures of adaptation in *Prochlorococcus* genomes

From community-level back to genome-level, we evaluated the pertinence of using fCUB to characterize the genomic diversity and ecology within a widely distributed and highly abundant group of marine prokaryotes, the genus *Prochlorococcus* (*51*). We first identified 261 *Prochlorococcus* MAGs within the MGnify marine genome catalogue and computed 123 fCUB in functions present in at least 85% of the MAGs. fCUBs were projected in a Principal Component Analysis (PCA) plot to visualize the diversity of functional signatures (Figure 4-A). It revealed an arch like structure with a continuum of variation across the signatures. We were not able to identify distinct groups via the PCA, and distributions of fCUB showed no multimodality, which would have favoured the idea of groups (Hartigan’s dip test for unimodality, p-values>0.05, n_bootstrap=10000). It is instead a progressive gradient of fCUB that forms along the continuum of individual species and ecotypes. This may be due to the fact that we only considered functions shared by the majority of the genus, i.e. core functions rather than accessory functions, essential for adaptation to specific niches (*52*). This is due to two factors: (i) we retain only functions found in the majority of *Prochlorococcus* MAGs to allow comparison of their fCUB, hence filtering accessory functions. And (ii) because accessory functions tend to be missed when reconstructing MAGs (*53*), limiting their numbers in our analysis. Thus, this framework does not capture all signals of ecological adaptation, but provides a tractable approach to characterize genomes and especially MAGs given the difficulty to reconstruct accessory genes in MAGs (*53*).

**Figure 4:**
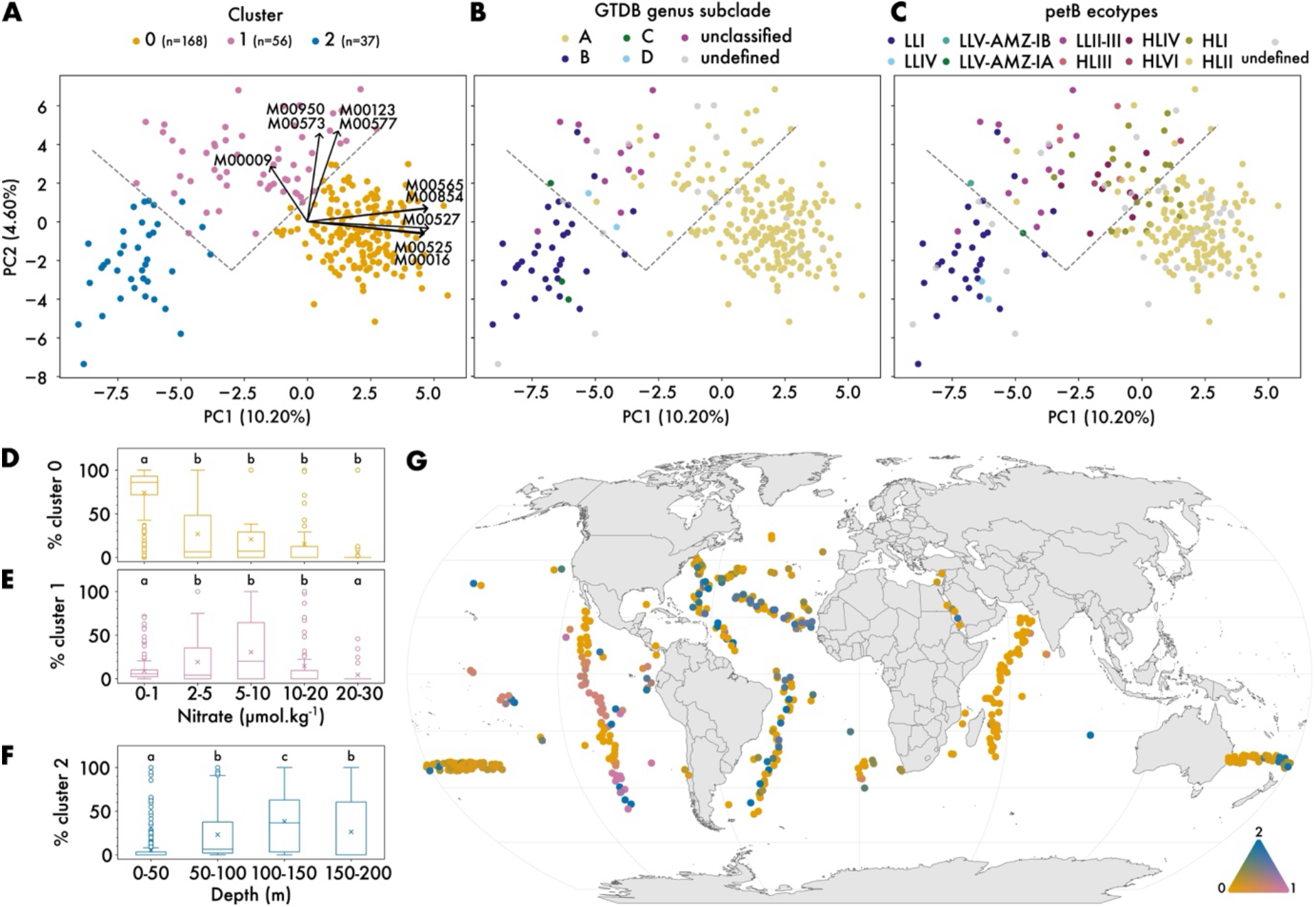
Functional bias within Prochlorococcus genus reveals genomic adaptation to ecological niches. **A**. Principal Component Analysis (PCA) biplot of 261 *Prochlorococcus* MAGs from Mgnify database characterized by 123 fCUB. Three clusters are identified with a K-means algorithm. Arrows originating from the point of coordinate (0,0) are the 5 most contributing functions of each PCA axis. The grey dashed line roughly separates clusters to facilitate the comparison with plot B and C. **B**. Same plot as A but coloured by GTDB taxonomy. The label “undefined” corresponds to MAGs of the *Prochlorococcus* genus without more information. The label “unclassified” corresponds to MAGs of the *Prochlorococcus* genus with more information but not belonging to any of A, B, C, D groups. **C**. Same plot as A and B but coloured by *petB* clades, used here as proxies for *Prochlorococcus* ecotypes (*50*). LL=low light, HL=high light. **D.E.F**. Proportion of a given cluster across ranges of nitrate concentration or depth. Crosses represent means. Different letters correspond to statistically significant different means (permutation test, n=10000, p-value<0.05). **G**. Mapping of the proportion of each cluster defined in A. Colours refers to the ternary diagram in the bottom right. If the colour is pink, that means that 100% of the *Prochlorococcus* in this community belongs to cluster 1. If colour is green, 50% of the *Prochlorococcus* found in this community belong to cluster 0, 50% to cluster 2, 0% to cluster 1. Note that a small jitter was added on the map to distinguish communities with similar coordinates but different depths.

We then applied a K-means clustering algorithm to identify three clusters along the functional bias continuum. The purpose of this clustering was to identify the main functions driving this spectrum, not to form groups. These three clusters are coherent with two independent taxonomic annotations: (i) using the Genome Taxon Database (GTDB) (*54*) taxonomic annotations based on 120 gene markers (Figure 4-B), or (ii) using *petB*, a high-resolution taxonomic marker gene for cyanobacteria ecotype delineation, absent from the GTDB 120 gene markers (Figure 4-C). All identified GTDB subclades or *petB* ecotypes are grouped together along the continuum, revealing the ability of fCUB to capture intra-genus taxonomic differences. For instance, low light (LL) and hight light (HL) ecotypes are found in separated regions of the continuum (Figure 4-C). These three clusters are also ecologically informative as they appear to be structured by distinct environmental drivers (Figure 4-DEF): cluster 0 is mostly found in nutrient-poor waters, cluster 1 in nutrient-rich waters, and cluster 2 in below surface euphotic waters (50-200m). Hence, the functional bias spectrum of *Prochlorococcus* MAGs appears to be mainly structured within these three ecological niches, with individual genomes showing progressive shift in signature from one cluster to another. Considering functions contributing to PC1 axis (Figure 4-A), most functions are related to lysine synthesis. High biases are found in surface nitrogen depleted waters (cluster 0) whereas low biases are found in deep nutrient rich waters (cluster2) (*55*). Here again, as proposed above, bias in lysine synthesis emerges as a potential key marker for nitrogen depletion. For PC2 axis (Figure 4-A), we found it was mainly driven by the synthesis of biotin (B7 vitamin). The last step of the B7 pathway uses an enzyme encompassing 6 iron atoms (*56*). Interestingly, we identified this step as the most biased of the pathway, especially in cluster 1 (Figure S13), a cluster found in known iron limited environments (*57*). Thus, biotin dependence on iron could explains why we are able to capture similar patterns to iron-depleted environments. By mapping the proportion of these clusters in each metagenomic community (Figure 4-G), we captured patterns similar to previous studies (e.g., nitrogen/iron limiting shifts along P18 transects (*57*)), reinforcing the potential of fCUB-based methods to capture intra-genus genomic diversity for adaptation to specific ecological niches. Overall, these results illustrate the power of a genome-level functional CUB approach to provide useful insights on the adaptation and ecology of a highly prevalent and abundant marine prokaryote.

## Conclusion

Marine microbial communities drive global biogeochemical cycles, yet identifying key metabolic functions for ecological adaptation and responses to environmental change has remained elusive. Here, we demonstrate that codon usage bias (CUB), a genomic marker of translational efficiency, captures functional adaptation to temperature and nutrient regimes across 589 ocean microbial communities. By extending CUB analysis beyond ribosomal genes to 320 functional gene groups, we uncovered four community types with distinct adaptation signatures that map to specific temperature-nutrient environments. This genome-centric approach offers a previously untapped lens into the functional ecology of uncultured microbes. Unlike traditional trait inference from taxonomy or metabarcoding, CUB provides a direct genomic proxy for potential growth rate and functional optimization, which is robust to community composition heterogeneity. Our findings validate CUB as a powerful tool to decode adaptive strategies in natural microbial communities, from nitrogen flux optimization in nutrient-poor waters to polyamine detoxification in warmer environments. Beyond the ocean, this framework is transferable to other ecosystems such as soil, gut, or host-associated microbiomes, and potentially to eukaryotes, opening new avenues to link genotypes to ecosystem functions. Future work integrating transcriptomic and proteomic data will bridge the gap between genomic potential and expressed phenotypes, while expanding CUB analyses to diverse environmental gradients (e.g., oxygen, trace metals) will refine our understanding of microbial adaptation in situ. As climate change reshapes ocean temperature and nutrient landscapes, our approach provides a predictive framework to anticipate microbial responses and their cascading effects on global biogeochemistry.

## Authors contributions

S.C. and C.G. conceptualized the study. A.B., L.G. and C.G. assembled and curated the data. C.G. and S.C. developed the methodology. C.G. wrote the code. C.G. and S.C wrote the manuscript with inputs and comments from all co-authors. F.J., M.A. L. G. and S.C. acquired funding.

## Code and Data availability

All code and associated data used to produce analyses and figures is available at DOI 10.5281/zenodo.22141364

## Material & Methods

### Metagenomes and Metagenome-Assembled Genomes (MAGs)

We gathered data from four major oceanographic campaigns: *Tara* Oceans, Bio-GO-SHIP and GeoTraces. 589 prokaryotic metagenomes were downloaded from the European Nucleotide Archive (for *Tara* Oceans sample accession, see Salazar et al. 2019 supplementary material (*58*), SRP278138 for Bio-GO-SHIP, SRP081823 for Geotraces). Corresponding metagenomic reads were then used to estimate the species abundance of Metagenome-Assembled Genomes (MAGs) in each community. Here, we used sylph (*59*) (v0.8.0) to estimate genome abundance of MAGs dereplicated at 95% average nucleotide identity (ANI) within the MGnify marine genomes catalogue (*60*) (v2.0 2024-05-10 update). This catalogue regroups 13k MAGs of 90 different phyla with notably KEGG Orthology (KO) functional annotations. The three main phyla are Pseudomonadota, Bacteroidota and Cyanobacteriota (respectively 50%, 17%, 4% of the catalogue)

### Environmental and contextual data

For *Tara Oceans*, we used the metadata provided in Salazar et al., 2019 (*58*). For GeoTraces, we gathered data from ipd2021_v2 (https://www.geotraces.org/geotraces-intermediate-data-product-2021/) and converted bad quality measurements (i.e. QV:SEADATANET value equal 3,4,9) to NAs.

For Bio-GO-SHIP, we assembled all metadata possible available and listed in Larkin et al., 2021 (*21*). We focused on CTD measured metadata and used the supplementary file of Larkin et al. (*21*) to identify the NCBI SRA stations IDs. For cruises P18, A13.5, I07N, I09N, NH14, AE1319 and BVAL46, we used a combination of depth and station number to match the NCBI station ids. For AMT28 we identified two shifts in station id based on discrepancies in coordinates between the supplementary file of Larkin et al. (*21*) and the AMT28 metadata file. After correction, we visually checked if date, latitude, longitude, depth and temperature were coherent between both files. As for GeoTraces metadata, we converted low-quality measurements (FLAG_W value equals 2,4,6 or measurement value equals −999) to NAs.

To be comparable across all expeditions, we converted AE1319, NH14 and *Tara* Oceans nutrients concentrations from µmol.L^−1^ to µmol.kg^−1^ using *in situ* measured density when available. When missing, we estimated density with the *density*.*rho* function from gsw (*61*) Python package using salinity, temperature and pressure. Doing so we obtained temperature, nutrients (nitrate, phosphate and silicate) and depth information across for 589 metagenomes. Many campaign-specific parameters (e.g., trace metals for Geotraces, chlorophyll *a* and carbon in *Tara* Oceans, eukaryotes cells count in Bio-GO-SHIP) could not be integrated since presenting too many missing values when combining the expeditions. Finally, only stations for which we had temperature, nitrate, silicate, phosphate and depth information were retained for the analyses. In addition, we limited our analyses to the global ocean euphotic zone (samples above 200m depth), which corresponded to 91% of the samples.

Longhurst biomes delimitation were extracted from Marineregion.org (*62*). We loaded the .shp file to R with the sf::st_read function from the sf package(*63*). Upwelling regions were extracted from Global Ecosystem Typology (https://global-ecosystems.org/explore/groups/M1.9). They were imported in R using the sf package(*63*), transformed to fit the Robinson map projection and grouped by the sf::st_boundary function. Then we computed the closest distance to an upwelling boundary using the sf::st_distance function. Closest distance to shore was computed in a similar approach, using the coastal border loaded from the ne_countries function of the naturalearth package(*64*).

### Codon Usage Bias (CUB) and ribosomal CUB (rCUB) computation

The genetic code, linking codons in DNA to amino acid in protein, is degenerated. A given amino acid can be encoded by multiple codons. We could expect equivalent frequencies between codons of a given amino acid, meaning that codons are chosen at random. However, this is not the case and in most genomes some codons are preferred than others (*11*). Codon Usage Bias (CUB) aims to quantify this codon preference within a given genome, that is to quantify how far from expected are codon frequencies in a given sequence of interest. Multiple methods and formula have been developed to compute CUB (*65*). Herein, we used the Measure Independent of Length and Composition formula (MILC) (*65*). MILC is one of the most robust H0 CUB quantification method and does not requires prior knowledge nor convergence toward a set of optimal codon as H1 method does (*66*).

The MILC approach focuses on codon frequencies to calculate the CUB. The observed frequency (f) of a codon is simply its occurrence (n) divided by the sum of the occurrences of all codons coding for the same amino acid. For example, we count 4 occurrences of the codon AAA and 6 occurrences of the codon AAG in a given sequence. AAA and AAG are the codons of the lysine (Lys) amino acid. Then, their frequencies are simply:

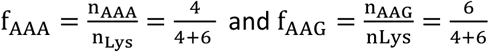

Observed frequencies can be computed at various biological organization levels (e.g., gene, pathway, genome, community) depending on the scope at which codons are counted. This observed frequency is then compared to the expected frequency of each codon. Expected frequency is sampled from the global composition of codons in a set of reference sequences relevant to the scope you are evaluating the bias for. For example, if you want to evaluate how biased is a gene in a pathway, you can consider the whole pathway as your relevant expected background.

In this study, we work at the genome-level scope and always compare the codons of genes (e.g., ribosomal genes) to the rest of their genome. The idea is to take into account the specificity of each genome (e.g., Guanine Cytosine rate, availability of transfer Ribonucleic Acid (tRNA)) and evaluate how different is a given group of gene to this background. In the event that a globally highly biased genome is taken as reference (i.e., very few codons are used), we are still able tell how much more biased is this group of genes compared to this reference. So here, each bias computed here is relative to each genome.

Observed (f_c_) and expected frequencies (g_c_) are compared using the MILC equation (*65*) that outputs a unique bias value aggregating the biases of all 64 codons:

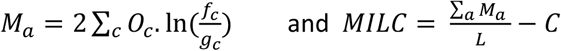

with *M*_*a*_ the measure of bias of a given amino acid. It sums the bias of all codons (c) encoding this amino acid. For each codon, we multiply the number of times the given codon is found in the gene (*O*_*c*_) by the natural logarithm of the *f*_*c*_ over *g*_*c*_ ratio. All *M*_*a*_ are summed to form the MILC. This sum is corrected by gene length (L) in number of codons and by the correction factor (C) which limits the mis-estimation of MILC in very long or very short genes, respectively (*65*).

The MILC formula outputs a CUB value per gene, for all genes within a given genome. To evaluate bias of a group of genes, we used the formula implemented in gRodon2 (*12, 67*). gRodon2 was built to study the CUB of genes encoding ribosomal proteins (rCUB) in order to predict growth rate. It computes the median MILC of all genes encoding ribosomal proteins, genes expected to be highly expressed (HE) during growth explaining why it is labelled as CUBHE. But we can actually study any group of genes using this approach (see below *fCUB*). In gRodon2, the median of MILC of all genes of the genome is named CUB. Differential CUB (dCUB) compares CUB and CUBHE given the following formula:

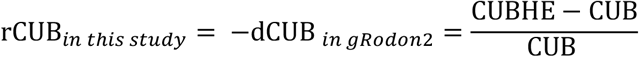

In short, dCUB is a genome-wide metric that compares the aggregated bias of all codons across all genes (CUB) versus the aggregated bias of all codons from genes of interest (CUBHE). To make rCUB easier to interpret, we computed opposite dCUB which corresponds to dCUB * (−1). Indeed, highly expressed genes tends to be more biased than the rest of the genome making the dCUB = (CUB-CUBHE)/CUB negative. By using the opposite dCUB, the metric becomes proportional to growth rate : the higher the rCUB the higher the maximum growth potential. Hence, over-biased ribosomal genes have positive rCUB and under-biased ribosomal genes (compared to the reference) have negative rCUB.

To simplify notations, we will refer to the concept of codon preference using the CUB acronym. In this study we label gRodon2’s -dCUB as rCUB to emphasize the focus on ribosomal genes and to distinguish it from bias of functional related group of genes, i.e. functional CUB (fCUB).

### Functional CUB (fCUB) computation

We generalized rCUB to other functional group of genes. Given CUB is a marker of protein dosage (*11*), we thought it was interesting to study it for protein-coding genes carrying other functions than ribosomal genes. We label it fCUB for **C**odon **U**sage **B**ias of **f**unctional related group of genes. To this aim, we detected KEGG modules (*11*) from MAG annotations. A module lists all the enzymes of the pathway allowing to transform a given substrate into final metabolite(s). Here, we computed the KEGG module bias using the same formula used to compute the rCUB, but replacing CUBHE by the median of the group of gene in the module of interest. A bias was computed only if the KEGG module was at least 50% complete in a given MAG. To compute module completeness, we used the kegg-pathways-completeness-tool (https://github.com/EBI-Metagenomics/kegg-pathways-completeness-tool) (*60*). As stated above, a positive fCUB reflects a group of gene more biased than the reference, and a negative fCUB a group of gene that is less biased compared to the reference.

### Quality filtering on MAGs

For most analyses, all MGnify MAGs were used without further filtering on quality (completeness/contamination) and thus corresponded to Medium-Quality (MQ) MAGs on completeness (>50%) and High-Quality (HQ) MAGs on contamination (<5%). This ensures low contamination to avoid polluting our codon frequencies with sequences likely originating from other genomes, and enough completeness to get sufficient genes in a given group of interest and sufficient genes to sample background codon composition.

For CUB computation, we only worked with MAGs above a given threshold when evaluating the bias of the group of interest. When computing rCUB, which focuses on ribosomal genes, we only worked with MAGs with 10 or more ‘ribosomal gene’ (91% of the 13k MAGs). When working with fCUB, we only computed a bias if at least 50% of a given functional group of gene was detected in the MAG (i.e., completion of functional KEGG module equal or above 50%).

Nevertheless, we also evaluated how our results in Figure 1 where sensible to quality threshold by looking only at HQ (completeness >90%, contamination <5%) MAGs. We reached similar conclusions (Figure S3-G)

### rCUB aggregation at community level: rCoB

To estimate the ribosomal biases at community level, we computed **r**ibosomal **Co**mmunity **B**ias (rCoB, i.e. rCUB at the community level) in a given metagenome *j* by multiplying each MAG rCUB (*rCUB*_*i*_) by its normalised abundance in the community (*γ*_*i*_). It is a weighted mean.

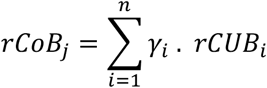

This is similar to the formula used by Abreu & Dal Bello to compute the MCN (*3*). It reveals the overall tendency of potential growth rate at the community level, led by high abundant and high biased MAGs.

### fCUB aggregation at community level: fCoB

For **f**unctional **Co**mmunity **B**ias (fCoB), we used exactly the same method as for rCoB.

fCoB in a given metagenome *j* is obtained by multiplying each MAG rCUB (*fCUB*_*i*_) by its re-normalised abundance (*γ*′_*i*_) of MAGs bearing the function *f* in the community. It is a weighted mean.

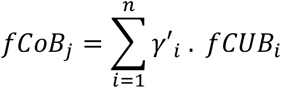

*γ*′_*i*_ is often different from *γ*_*i*_. Indeed, usually a function is not bearded by all MAGs of a community. We remove abundances of non-bearing MAGs, renormalize to sum to 1 and use these *γ*′_*i*_ abundances in fCoB. Hence, the community bias reflects the mean fCUB of a randomly sampled cell containing the function, and not of a randomly sampled cell of the community (interpretation adapted from (*3*)). It is mainly influenced by high abundant function bearer MAGs.

At the end, we obtain a fCoB for each function detected at least in one genome of the community. Most function were quasi-universal (see main text) meaning that they were detected in most metagenomes of our study. When a function was not detected in a given community, we artificially set its value to 0. The 320 selected functions are at least detected in 50% of our communities (i.e. 50% universality).

### fCoB ordination analysis (UMAP)

The dataset was scaled, and projected using an ordination algorithm, here Uniform Manifold Approximate Projection (UMAP (*68*)). From the 589 metagenomes gathered in this study, we kept only the 485 from the 0 to 100 meters depth for the UMAP analysis for two main reasons. First, we identified that most of the below 100 meters depth metagenomes were coming from a single region of the globe, namely South Atlantic, hence making hard to decipher between an Atlantic signal and a depth signal. Second, when plotting the UMAP while keeping these points between 100 and 200 meters, a cluster of deep metagenomes formed. However, because of its divergence to the other communities, keeping these deep metagenomes worsen dispersion of the other UMAP clusters (Figure S11), hence complicating interpretability.

We also evaluated how setting missing values to 0 was impacting the UMAP. Indeed, it’s a trade-off: we could think of keeping only 100% universal function, but doing so we could exclude very informative functions with only few missing values. On the other hand, including a function with a lot of missing value could artificially constrains our UMAP by grouping lots of zero imputed with zero natural values. To evaluate this impact, we plotted a UMAP by keeping only function with at least 50%, 85%, 95% and 100% coverage. We were still able to identify the same clusters even if their limits faded (Figure S8).

This likely means there is still enough statistical power to form groups using very universal functions, from central housekeeping metabolism, but it also suggests that less abundant functions are important to delimit the clusters. For the UMAP presented in Figure 3, we worked with function found in at least 50% of our communities, i.e. 50 % universality, selecting 320 functions.

### Adaptive functional signatures in temperature-nutrients regimes

For identifying functions driving the distinct clusters on the UMAP, we first visually defined boarders of each cluster. We run an extreme gradient boosting algorithm using the python library XGBoost (*40*) to identify function helpful to distinguish one cluster from the three others. It is inspired by other UMAP cluster analysis methods (*69*). Functions are ordered by total gain which measure how important a given function was to segregate a given cluster from the three others during training. For XGBoost training, we selected only function with universality above 90 % and function not showing high global correlation (Figure 2, Figure S7), in order to focus on modules with local pattern. We also removed very extreme values of biases (replaced by 0) in order to limit learning on outliers not representative of the cluster. Extreme values were defined relatively to inter-quartile range (IQR=Q3-Q1):

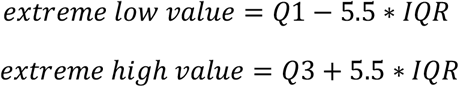

Among these high total gain function, we selected three representatives (Figure S12). We made sure to select positively and negatively biased typical functions for each cluster, to show that both can be selected in every cluster. We do not aim to provide an exhaustive characterisation of each cluster specificity, but rather to provide insights on some functions driving them.

### MAGs contribution to temperature vs. rCoB correlation

MAG contribution was explored in order to identify if the correlation between temperature and rCoB was under the influence of some specific prokaryotes, or on the contrary based on a signal spread in the community. To do so, we defined contribution of an individual *i* (MAG for rCUB or ASV for MCN) as the ratio between its term in the community mean and the value of the community mean. So it is the abundance (*γ*_*i*_) time the metric (*rCUB*_*i*_ or *n*_16*S*_*i*_, the number of 16S rRNA genes in *ASV*_*i*_) of a given individual divided by the averaged value (*rCoB*_*c*_ or *MCN*_*c*_(*3*)) of the community *c* it was detected in.

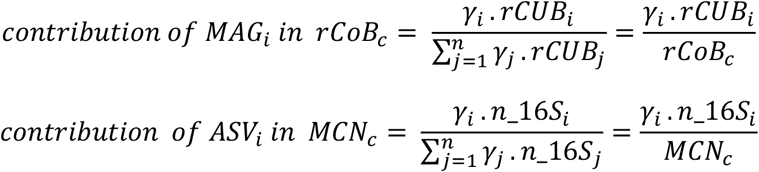

Individual with high contribution have a strong influence on their rCoB. When rCUB was negative (it can happen if there is less optimisation in ribosomal genes compared to the background composition), we compared the absolute value of the individual to the absolute value of the community, hence capturing a contribution effect. Those individuals were labelled as “negative contributors”. We then summed the contribution of each individual across all stations. If the correlation temperature versus metric (rCoB or MCN) is mainly explained by some specific individuals, the slope of this correlation should flatten after removing those individuals. To verify this, we progressively removed individuals in our experiment, starting with individuals with the biggest contribution across stations. The plots in Figure S3-AB represent the evolution of this slope while removing progressively the highest contributors.

### Statistical models

To understand the relationship between temperature, nutrient concentration and rCoB, we explored various linear models using R (version 4.4.1) lm function.

First, we tested the relationship between distance to shore (continent or island) and distance to upwelling region (see computation above in Material & Methods). We constructed three models:

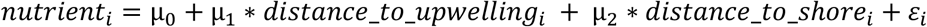

*nutrient*_*i*_ is successively one of the three nutrients studied: nitrate, phosphate or silicate concentrations of metagenome *i* and *ε*_*i*_ the residuals.

Then, we tested the relationship between rCoB and environmental parameters.

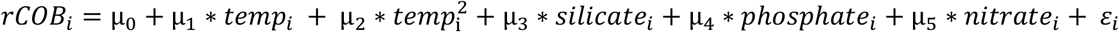

*temp* and *temp*^*2*^ *a*re the two components of the polynomial of order two regression between temperature and rCoB found in Figure 1. *i* is the indice of the given station and *ε*_*i*_ the residuals. Note that µ_0_ integrate the c constant of the equation linking rCoB to temperature in a polynomial of order 2 fit:

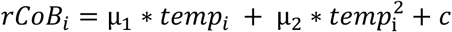

The normality of the residuals for all models was assessed with a Shapiro test.

Following the study of the model outputs, we performed a contribution analysis using the Lindeman, Merenda, and Gold (LMG) parameter of the calc.relimp function in the relaimpo R package (*25*). The aim of this approach is to extract the contribution of variables to the total variability of the model. Knowing that the order at which variables enter a linear model can affect their contribution, the LMG approach average the contribution across all permutations of variable order.

We computed the LMG contribution of temperature and nutrient by group: *temp* and *temp*^*2*^ under the temperature label and *silicate, phosphate and nitrate* under the nutrient label. Grouping allows to assess the contribution of variables as a block, limiting overlap between correlated metrics. It is useful to study the nutrient effect composed of three variables (*silicate, phosphate, nitrate*) versus the temperature effect composed of two variables (*temp* and *temp*^*2*^).

### *Prochlorococcus* intra-genus CUB analysis

We extracted 263 MAGs MQ-HQ assigned to the genus *Prochlorococcus* from MGnify. We followed the same procedure described above to compute functional biases fCUB. We removed all functions not detected in at least in 85% of the MAGs. Missing values were set to zero and a Principal Component Analysis (PCA) ordination was performed using the scikit-learn python package (*70*). Two MAGs exhibited markedly different fCUB signatures and were clear outliers in the PCA, potentially reflecting taxonomic mislabelling. These two MAGs were removed from the analysis (MGYG000476853 and MGYG00460174) to facilitate visualisation of the remaining MAGs. Based on this PCA, three clusters were identified using K-means clustering (Figure 4-BEFG) (*70*). Because our aim was not to define groups with meaningful limits, but rather to characterize steps along a continuum, we did not determine the number of clusters using the classical inertia drop analysis. Instead, we arbitrary selected three clusters to facilitate interpretation of the ternary diagram shown in Figure 4-G.

These clusters were also ecologically relevant, as shown by the environmental range analysis that followed. For this analysis, we computed the proportion of each cluster within each community and evaluated its enrichment across ranges of nutrient concentrations and depth.

## Bibliography

1. C. B. Field, M. J. Behrenfeld, J. T. Randerson, P. Falkowski, Primary Production of the Biosphere: Integrating Terrestrial and Oceanic Components. Science 281, 237–240 (1998).

2. E. J. Zakem, J. McNichol, J. L. Weissman, Y. Raut, L. Xu, E. R. Halewood, C. A. Carlson, S. Dutkiewicz, J. A. Fuhrman, N. M. Levine, Functional biogeography of marine microbial heterotrophs. Science 388, eado5323 (2025).

3. C. I. Abreu, M. Dal Bello, C. Bunse, J. Pinhassi, J. Gore, Warmer temperatures favor slower-growing bacteria in natural marine communities. Science Advances 9, eade8352 (2023).

4. J. Ramoneda, E. Stallard-Olivera, M. Hoffert, C. C. Winfrey, M. Stadler, J. P. Niño-García, N. Fierer, Building a genome-based understanding of bacterial pH preferences. Science Advances 9, eadf8998 (2023).

5. S. Louca, L. W. Parfrey, M. Doebeli, Decoupling function and taxonomy in the global ocean microbiome. Science 353, 1272–1277 (2016).

6. S. Raguideau, S. Plancade, N. Pons, M. Leclerc, B. Laroche, Inferring Aggregated Functional Traits from Metagenomic Data Using Constrained Non-negative Matrix Factorization: Application to Fiber Degradation in the Human Gut Microbiota. PLOS Computational Biology 12, e1005252 (2016).

7. N. Fierer, A. Barberán, D. C. Laughlin, Seeing the forest for the genes: using metagenomics to infer the aggregated traits of microbial communities. Front. Microbiol. 5 (2014).

8. Z. Li, W. J. Riley, G. L. Marschmann, U. Karaoz, I. A. Shirley, Q. Wu, N. J. Bouskill, K.-Y. Chang, P. M. Crill, R. F. Grant, E. King, S. R. Saleska, M. B. Sullivan, J. Tang, R. K. Varner, B. J. Woodcroft, K. C. Wrighton, E. L. Brodie, A framework for integrating genomics, microbial traits, and ecosystem biogeochemistry. Nat Commun 16, 2186 (2025).

9. J. S. Madin, D. A. Nielsen, M. Brbic, R. Corkrey, D. Danko, K. Edwards, M. K. M. Engqvist, N. Fierer, J. L. Geoghegan, M. Gillings, N. C. Kyrpides, E. Litchman, C. E. Mason, L. Moore, S. L. Nielsen, I. T. Paulsen, N. D. Price, T. B. K. Reddy, M. A. Richards, E. P. C. Rocha, T. M. Schmidt, H. Shaaban, M. Shukla, F. Supek, S. G. Tetu, S. Vieira-Silva, A. R. Wattam, D. A. Westfall, M. Westoby, A synthesis of bacterial and archaeal phenotypic trait data. Sci Data 7, 170 (2020).

10. S. Vieira-Silva, E. P. C. Rocha, The Systemic Imprint of Growth and Its Uses in Ecological (Meta)Genomics. PLOS Genetics 6, e1000808 (2010).

11. T. Ikemura, Codon usage and tRNA content in unicellular and multicellular organisms. Molecular Biology and Evolution, doi: 10.1093/oxfordjournals.molbev.a040335 (1985).

12. J. L. Weissman, S. Hou, J. A. Fuhrman, Estimating maximal microbial growth rates from cultures, metagenomes, and single cells via codon usage patterns. Proceedings of the National Academy of Sciences 118, e2016810118 (2021).

13. G. Boël, R. Letso, H. Neely, W. N. Price, K.-H. Wong, M. Su, J. D. Luff, M. Valecha, J. K. Everett, T. B. Acton, R. Xiao, G. T. Montelione, D. P. Aalberts, J. F. Hunt, Codon influence on protein expression in E. coli correlates with mRNA levels. Nature 529, 358–363 (2016).

14. F. Hia, S. F. Yang, Y. Shichino, M. Yoshinaga, Y. Murakawa, A. Vandenbon, A. Fukao, T. Fujiwara, M. Landthaler, T. Natsume, S. Adachi, S. Iwasaki, O. Takeuchi, Codon bias confers stability to human mRNAs. EMBO Rep 20, e48220 (2019).

15. G. Hanson, J. Coller, Codon optimality, bias and usage in translation and mRNA decay. Nat Rev Mol Cell Biol 19, 20–30 (2018).

16. W. D. Chang, Y.-J. Choe, Quality control and signaling pathways at stalled ribosomes. Exp Mol Med 58, 82–93 (2026).

17. G. Hanson, J. Coller, Codon optimality, bias and usage in translation and mRNA decay. Nat Rev Mol Cell Biol 19, 20–30 (2018).

18. M. Roller, V. Lucić, I. Nagy, T. Perica, K. Vlahoviček, Environmental shaping of codon usage and functional adaptation across microbial communities. Nucleic Acids Res 41, 8842–8852 (2013).

19. T. Antman, O. Lewin-Epstein, T. Yerushalmi, Y. S. Broder, D. Zeevi, Global microbial DNA signatures of temperature and nutrient limitation across ecosystems. Nat Microbiol, 1–11 (2026).

20. S. Sunagawa, S. G. Acinas, P. Bork, C. Bowler, Tara Oceans Coordinators, D. Eveillard, G. Gorsky, L. Guidi, D. Iudicone, E. Karsenti, F. Lombard, H. Ogata, S. Pesant, M. B. Sullivan, P. Wincker, C. de Vargas, Tara Oceans: towards global ocean ecosystems biology. Nat Rev Microbiol 18, 428–445 (2020).

21. A. A. Larkin, C. A. Garcia, N. Garcia, M. L. Brock, J. A. Lee, L. J. Ustick, L. Barbero, B. R. Carter, R. E. Sonnerup, L. D. Talley, G. A. Tarran, D. L. Volkov, A. C. Martiny, High spatial resolution global ocean metagenomes from Bio-GO-SHIP repeat hydrography transects. Sci Data 8, 107 (2021).

22. S. J. Biller, P. M. Berube, K. Dooley, M. Williams, B. M. Satinsky, T. Hackl, S. L. Hogle, A. Coe, K. Bergauer, H. A. Bouman, T. J. Browning, D. De Corte, C. Hassler, D. Hulston, J. E. Jacquot, E. W. Maas, T. Reinthaler, E. Sintes, T. Yokokawa, S. W. Chisholm, Marine microbial metagenomes sampled across space and time. Sci Data 5, 180176 (2018).

23. C. A. Garcia, G. I. Hagstrom, A. A. Larkin, L. J. Ustick, S. A. Levin, M. W. Lomas, A. C. Martiny, Linking regional shifts in microbial genome adaptation with surface ocean biogeochemistry. Philos Trans R Soc Lond B Biol Sci 375, 20190254 (2020).

24. A. R. Longhurst, Ecological Geography of the Sea (San Diego [u.a.] Acad. Press, 1998; http://archive.org/details/ecologicalgeogra01long).

25. U. Grömping, Relative Importance for Linear Regression in R: The Package relaimpo. Journal of Statistical Software 17, 1–27 (2006).

26. A. W. Visser, P. Brun, S. Chakraborty, T. S. Dencker, P. D. van Denderen, R. van Gemert, H. van Someren Gréve, I. Heilmann, M. W. Holm, S. H. Jónasdóttir, K. M. Kenitz, T. Kiørboe, M. Lindegren, P. Mariani, L. T. Nielsen, M. Pancic, M. Payne, L. Pécuchet, N. A. Schnedler-Meyer, U. H. Thygesen, A. Törnroos, K. H. Andersen, Seasonal strategies in the world’s oceans. Progress in Oceanography 189, 102466 (2020).

27. A. Burga, B. Lehner, Beyond genotype to phenotype: why the phenotype of an individual cannot always be predicted from their genome sequence and the environment that they experience. The FEBS Journal 279, 3765–3775 (2012).

28. D. L. Kirchman, X. A. G. Morán, H. Ducklow, Microbial growth in the polar oceans — role of temperature and potential impact of climate change. Nat Rev Microbiol 7, 451–459 (2009).

29. J. L. Weissman, A. Walling, H. Ducklow, E. J. Zakem, Genomic traits associated with copiotrophy decouple from maximum growth rate predictions along temperature gradients. ISME J 20, wrag147 (2026).

30. M. Kanehisa, M. Furumichi, Y. Sato, Y. Matsuura, M. Ishiguro-Watanabe, KEGG: biological systems database as a model of the real world. Nucleic Acids Res 53, D672–D677 (2025).

31. J. L. López, A. Litos, M. Siemers, M. D. Boer, E. Padial, P. Doijad, B. E. Dutilh, Widespread synchronization of codon usage in functionally related genes.

32. S. Kurihara, S. Oda, Y. Tsuboi, H. Kim, M. Oshida, H. Kumagai, H. Suzuki,- Glutamylputrescine Synthetase in the Putrescine Utilization Pathway of Escherichia coli K-12. The Journal of biological chemistry 283, 19981–90 (2008).

33. S. Sugiyama, S. Ishikawa, H. Tomitori, M. Niiyama, M. Hirose, Y. Miyazaki, K. Higashi, M. Murata, H. Adachi, K. Takano, S. Murakami, T. Inoue, Y. Mori, K. Kashiwagi, K. Igarashi, H. Matsumura, Molecular mechanism underlying promiscuous polyamine recognition by spermidine acetyltransferase. The International Journal of Biochemistry & Cell Biology 76, 87–97 (2016).

34. K. Igarashi, K. Kashiwagi, Effects of polyamines on protein synthesis and growth of Escherichia coli. J Biol Chem 293, 18702–18709 (2018).

35. S. Muduli, S. Karmakar, S. Mishra, The coordinated action of the enzymes in the L-lysine biosynthetic pathway and how to inhibit it for antibiotic targets. Biochimica et Biophysica Acta (BBA) - General Subjects 1867, 130320 (2023).

36. O. Irazoki, S. B. Hernandez, F. Cava, Peptidoglycan Muropeptides: Release, Perception, and Functions as Signaling Molecules. Front. Microbiol. 10 (2019).

37. H. Akashi, T. Gojobori, Metabolic efficiency and amino acid composition in the proteomes of Escherichia coli and Bacillus subtilis. Proc Natl Acad Sci U S A 99, 3695–3700 (2002).

38. J. Kleinteich, F. Hildebrand, M. Bahram, A. Y. Voigt, S. A. Wood, A. D. Jungblut, F. C. Küpper, A. Quesada, A. Camacho, D. A. Pearce, P. Convey, W. F. Vincent, C. Zarfl, P. Bork, D. R. Dietrich, Pole-to-Pole Connections: Similarities between Arctic and Antarctic Microbiomes and Their Vulnerability to Environmental Change. Front. Ecol. Evol. 5 (2017).

39. S. Chaffron, E. Delage, M. Budinich, D. Vintache, N. Henry, C. Nef, M. Ardyna, A. Zayed, P. Junger, P. Galand, C. Lovejoy, A. Murray, H. Sarmento, S. Acinas, M. Babin, D. Iudicone, O. Jaillon, E. Karsenti, P. Wincker, D. Eveillard, Environmental vulnerability of the global ocean epipelagic plankton community interactome. Science Advances 7, eabg1921 (2021).

40. T. Chen, C. Guestrin, XGBoost: A Scalable Tree Boosting System. Proceedings of the 22nd ACM SIGKDD International Conference on Knowledge Discovery and Data Mining, 785–794 (2016).

41. J. C. Kaiser, D. E. Heinrichs, Branching Out: Alterations in Bacterial Physiology and Virulence Due to Branched-Chain Amino Acid Deprivation. mBio 9, 10.1128/mbio.01188-18 (2018).

42. M. Jespersen, T. Wagner, Assimilatory sulfate reduction in the marine methanogen Methanothermococcus thermolithotrophicus. Nat Microbiol 8, 1227–1239 (2023).

43. T. Weber, N. A. Wiseman, A. Kock, Global ocean methane emissions dominated by shallow coastal waters. Nat Commun 10, 4584 (2019).

44. P. Santucci, M. D. Johansen, V. Point, I. Poncin, A. Viljoen, J.-F. Cavalier, L. Kremer, S. Canaan, Nitrogen deprivation induces triacylglycerol accumulation, drug tolerance and hypervirulence in mycobacteria. Sci Rep 9, 8667 (2019).

45. C. M. Callbeck, D. E. Canfield, M. M. M. Kuypers, P. Yilmaz, G. Lavik, B. Thamdrup, C. J. Schubert, L. A. Bristow, Sulfur cycling in oceanic oxygen minimum zones. Limnology and Oceanography 66, 2360–2392 (2021).

46. C. De Falco, F. Desbiolles, A. Bracco, C. Pasquero, Island Mass Effect: A Review of Oceanic Physical Processes. Front. Mar. Sci. 9 (2022).

47. I. Kushkevych, V. Procházka, M. Vítězová, D. Dordević, M. Abd El-Salam, S. K.-M. R. Rittmann, Anoxygenic photosynthesis with emphasis on green sulfur bacteria and a perspective for hydrogen sulfide detoxification of anoxic environments. Front Microbiol 15, 1417714 (2024).

48. S. Neukirchen, I. A. C. Pereira, F. L. Sousa, Stepwise pathway for early evolutionary assembly of dissimilatory sulfite and sulfate reduction. ISME J 17, 1680–1692 (2023).

49. A. Sekowska, H. Ashida, A. Danchin, Revisiting the methionine salvage pathway and its paralogues. Microbial Biotechnology 12, 77–97 (2019).

50. G. K. Farrant, H. Doré, F. M. Cornejo-Castillo, F. Partensky, M. Ratin, M. Ostrowski, F. D. Pitt, P. Wincker, D. J. Scanlan, D. Iudicone, S. G. Acinas, L. Garczarek, Delineating ecologically significant taxonomic units from global patterns of marine picocyanobacteria. Proceedings of the National Academy of Sciences 113, E3365–E3374 (2016).

51. F. Partensky, L. Garczarek, Prochlorococcus: Advantages and Limits of Minimalism. Annual Review of Marine Science 2, 305–331 (2010).

52. D. Medini, C. Donati, H. Tettelin, V. Masignani, R. Rappuoli, The microbial pan-genome. Current Opinion in Genetics & Development 15, 589–594 (2005).

53. T. A. Pellegrinetti, J. Molligan, L. W. Mendes, A. Pedrinho, E. Pérez-López, Rethinking metagenome-assembled genome completeness: are we truly recovering complete genomes? Front. Microbiol. 17 (2026).

54. P.-A. Chaumeil, A. J. Mussig, P. Hugenholtz, D. H. Parks, GTDB-Tk: a toolkit to classify genomes with the Genome Taxonomy Database. Bioinformatics 36, 1925–1927 (2020).

55. F. Partensky, L. Garczarek, Prochlorococcus: Advantages and Limits of Minimalism. Annual Review of Marine Science 2, 305–331 (2010).

56. F. Berkovitch, Y. Nicolet, J. T. Wan, J. T. Jarrett, C. L. Drennan, Crystal Structure of Biotin Synthase, an S-Adenosylmethionine-Dependent Radical Enzyme. Science 303, 76–79 (2004).

57. L. J. Ustick, A. A. Larkin, C. A. Garcia, N. S. Garcia, M. L. Brock, J. A. Lee, N. A. Wiseman, J. K. Moore, A. C. Martiny, Metagenomic analysis reveals global-scale patterns of ocean nutrient limitation. Science 372, 287–291 (2021).

58. G. Salazar, L. Paoli, A. Alberti, J. Huerta-Cepas, H.-J. Ruscheweyh, M. Cuenca, C. M. Field, L. P. Coelho, C. Cruaud, S. Engelen, A. C. Gregory, K. Labadie, C. Marec, E. Pelletier, M. Royo-Llonch, S. Roux, P. Sánchez, H. Uehara, A. A. Zayed, G. Zeller, M. Carmichael, C. Dimier, J. Ferland, S. Kandels, M. Picheral, S. Pisarev, J. Poulain, S. G. Acinas, M. Babin, P. Bork, E. Boss, C. Bowler, G. Cochrane, C. de Vargas, M. Follows, G. Gorsky, N. Grimsley, L. Guidi, P. Hingamp, D. Iudicone, O. Jaillon, S. Kandels-Lewis, L. Karp-Boss, E. Karsenti, F. Not, H. Ogata, S. Pesant, N. Poulton, J. Raes, C. Sardet, S. Speich, L. Stemmann, M. B. Sullivan, S. Sunagawa, P. Wincker, S. G. Acinas, M. Babin, P. Bork, C. Bowler, C. de Vargas, L. Guidi, P. Hingamp, D. Iudicone, L. Karp-Boss, E. Karsenti, H. Ogata, S. Pesant, S. Speich, M. B. Sullivan, P. Wincker, S. Sunagawa, Gene Expression Changes and Community Turnover Differentially Shape the Global Ocean Metatranscriptome. Cell 179, 1068–1083.e21 (2019).

59. J. Shaw, Y. W. Yu, Rapid species-level metagenome profiling and containment estimation with sylph. Nat Biotechnol, 1–12 (2024).

60. L. Richardson, B. Allen, G. Baldi, M. Beracochea, M. L. Bileschi, T. Burdett, J. Burgin, J. Caballero-Pérez, G. Cochrane, L. J. Colwell, T. Curtis, A. Escobar-Zepeda, T. A. Gurbich, V. Kale, A. Korobeynikov, S. Raj, A. B. Rogers, E. Sakharova, S. Sanchez, D. J. Wilkinson, R. D. Finn, MGnify: the microbiome sequence data analysis resource in 2023. Nucleic Acids Res 51, D753–D759 (2023).

61. E. Firing, Filipe, A. Barna, R. Abernathey, TEOS-10/GSW-Python: v3.4.1.post0, Zenodo (2021); 10.5281/zenodo.5214122.

62. Flanders Marine Institute, Longhurst Provinces (2009). https://www.marineregions.org/sources.php#longhurst.

63. E. Pebesma, R. Bivand, E. Racine, M. Sumner, I. Cook, T. Keitt, R. Lovelace, H. Wickham, J. Ooms, K. Müller, T. L. Pedersen, D. Baston, D. Dunnington, sf: Simple Features for R, version 1. 0–24 (2026); https://cran.r-project.org/web/packages/sf/index.html.

64. P. Massicotte, A. South, K. Hufkens, rnaturalearth: World Map Data from Natural Earth, version 1.2.0 (2026); https://cran.r-project.org/web/packages/rnaturalearth/index.html.

65. F. Supek, K. Vlahoviček, Comparison of codon usage measures and their applicability in prediction of microbial gene expressivity. BMC Bioinformatics 6, 182 (2005).

66. R. Merkl, A Survey of Codon and Amino Acid Frequency Bias in Microbial Genomes Focusing on Translational Efficiency. J Mol Evol 57, 453–466 (2003).

67. J. L. Weissman, M. Peras, T. P. Barnum, J. A. Fuhrman, Benchmarking Community-Wide Estimates of Growth Potential from Metagenomes Using Codon Usage Statistics. mSystems 7, e00745–22 (2022).

68. L. McInnes, J. Healy, N. Saul, L. Großberger, UMAP: Uniform Manifold Approximation and Projection. Journal of Open Source Software 3, 861 (2018).

69. P. Castellano-Escuder, D. K. Zachman, K. Han, M. D. Hirschey, GAUDI: interpretable multi-omics integration with UMAP embeddings and density-based clustering. Nat Commun 16, 5771 (2025).

70. F. Pedregosa, G. Varoquaux, A. Gramfort, V. Michel, B. Thirion, O. Grisel, M. Blondel, P. Prettenhofer, R. Weiss, V. Dubourg, J. Vanderplas, A. Passos, D. Cournapeau, M. Brucher, M. Perrot, É. Duchesnay, Scikit-learn: Machine Learning in Python. J. Mach. Learn. Res. 12, 2825–2830 (2011).

71. J. F. Caddy, A. Bakun, A tentative classification of coastal marine ecosystems based on dominant processes of nutrient supply. Ocean & Coastal Management 23, 201–211 (1994).

72. M. E. Taboada, P. C. Hernández, H. R. Galleguillos, E. K. Flores, T. A. Graber, Behavior of sodium nitrate and caliche mineral in seawater: Solubility and physicochemical properties at different temperatures and concentrations. Hydrometallurgy 113–114, 160–166 (2012).

73. H.-C. Cheon, S.-Y. Nam, S.-H. Kim, Effects of pH, Temperature, and Dissolved Oxygen on Phosphorus Release from Marine Sediment to Seawater. Journal of Navigation and Port Research 36, 513–519 (2012).

74. G. Okamoto, T. Okura, K. Goto, Properties of silica in water. Geochimica et Cosmochimica Acta 12, 123–132 (1957).

75. M. Ardyna, K. R. Arrigo, Phytoplankton dynamics in a changing Arctic Ocean. Nat. Clim. Chang. 10, 892–903 (2020).

76. J. I. Arroyo, P. A. Marquet, C. P. Kempes, G. West, Toward a General Theory for the Scaling and Universality of Thermal Responses in Biology. arXiv arXiv:2503.05128 [Preprint] (2025). 10.48550/arXiv.2503.05128.

77. M. K. Thomas, M. Aranguren-Gassis, C. T. Kremer, M. R. Gould, K. Anderson, C. A. Klausmeier, E. Litchman, Temperature–nutrient interactions exacerbate sensitivity to warming in phytoplankton. Global Change Biology 23, 3269–3280 (2017).

78. J. J. Elser, L. B. Stabler, R. P. Hassett, Nutrient limitation of bacterial growth and rates of bacterivory in lakes and oceans: a comparative study. Aquatic Microbial Ecology 09, 105–110 (1995).

