## Supplementary Material for "Codon usage bias maps ocean microbiome adaptation to temperature-nutrients regimes"

### Supplementary text:

#### Text S1: Comparison between MCN and rCoB

Mean Copy Number (MCN) is an approach developed by Abreu and Dal Bello using metabarcoding detection of a bacteria followed by a taxonomic assignation of 16S RNA operon copy number(*3*) . A large MCN indicates greater relative abundance of fast-growing bacteria, while more abundant slow-growing bacteria drive the MCN to smaller values. rCoB correlates more with temperature than MCN, even when tweaking MCN to improve its correlation (Figure S3, CD). Indeed, we showed that limiting taxonomic match at the genus level was improving the MCN x temperature correlation, maybe because lowering the trait variability due to high matches. rCoB correlation with temperature was also more stable to highly contributing bacteria removal, revealing a signal more spread out in the community and less dependent on specific genus (Figure S3, AB). It might be explained by the continuous character of rCUB compared to the discrete character of 16S RNA operon copy number and the fact that no taxonomic assignment is needed for rCUB.

#### Text S2: Exploring the causal relationships of the negative correlation linking temperature to nutrient concentration in the global ocean

The observed negative association between nutrients concentration and temperature confounds the causal understanding of environmental factors constraining potential growth rate. To tackle this issue, we tested several causal hypotheses informed by previous knowledge. We tried to resolve the collinearity issues between temperature and nutrients, by exploring causes of nutrients distributions. We hypothesised that nitrate, phosphate and silicate distributions in the global ocean were explained by their proximity to coastal sources (continent, island or upwelling zone). The hypothesis is that those regions should harbour high nutrients concentrations due to coastal runoff, island mass effect and/or nutrient resuspension (*46*, *71*). We constructed basic linear models to explore this hypothesis (Material & Methods). In most models, the upwelling and coastal proximity were significantly explaining nutrient distributions. But those effects were very small and did not explain most of the variability (r2 nitrate=0.04, r2 phosphate =0.05, r2 silicate=0.02), which lead us to discard this hypothesis. It is probably explained by currents, with trajectories too different from straight lines. Hence at the global scale, and given the fact that most of the communities assembled in this study are far from coastal sources, distance to coast does not explain nutrient concentration.

When exploring other hypothesis to explain nutrient distribution, we looked at the negative relationship between temperature and nutrients ($r^2_{NO3}$=0.43 $\rho_{NO3}$=-0.66; $r^2_{PO4}$=0.39 $\rho_{PO4}$=-0.62; $r^2_{SiO4}$=0.19 $\rho_{SiO4}$=-0.43). It is known that higher temperatures increase solubility of dissolved ions (*72*–*74*), implying a positive relationship between nutrients and temperature. The fact we observe a negative relationship might be explained by another reason developed below.

Nutrients originally coming from coastal sources are spread by oceanic currents in the global ocean. Their trajectory differs from straight lines hence not linking nutrient distributions to straight shore nor upwelling distance. The negative correlation between temperature and nutrient concentrations would arise from two complementary processes. As temperature increases, the water column becomes more strongly stratified: warm, less dense surface waters mix less effectively with the cooler, nutrient-rich deeper layers. At the same time, photosynthetic organisms rapidly consume the available nutrients in the warm, sunlit surface layer, further reducing their concentrations. Thus, we hypothesise that vertical thermal stratification enhanced by photosynthetic activity is the causal explanation of the negative relationship between nutrients and temperature in our dataset. This hypothesis does not include any autotrophy like nitrogen autotrophy capturing atmospheric nitrogen.

We did not include primary production in our models even if Kirchman et al. reported the high correlation between effective bacterial growth rate and primary production in the global ocean (*28*). In a causal framework, we could explain this link by the fact that high primary production creates a prosperous growing environment for heterotrophic bacteria by releasing labile dissolved organic molecules. We did not integrate any proxy of primary production in our causal framework since it was not measured for all 589 stations. Nevertheless, we know that nutrient availability is one control of primary production (*75*). Considering the fact the we controlled for nutrient effect on rCoB, we simultaneously controlled for the nutrient-originating primary production effect. Doing so, we cannot estimate the direct nutrient effect on rCoB, but only its total effect encompassing primary production, which is still informative.

### Text S3: Effect of temperature and nutrient on rCoB per Longhurst Biome

Given the fact that stratification enhanced by photosynthetic activity is not a cofounder of temperature and nutrient but a mediator (temperature ⟶ stratification ⟶ nutrient concentration), we constructed a multiple linear regression model explaining the distribution of rCoB by temperature and nutrients. The main hypothesis we tested is that rCoB distribution is explained by a combination of temperature and nutrient concentrations, that is to say by different temperature-nutrient regimes. Given it has been previously shown that temperature affects effective growth rates (*76*) (phenotypic growth rate), we hypothesized it could impact genome-encoded potential growth rates (genotypic growth rate) and might be captured at the community level by rCoB. We also considered the same hypothesis linking nutrient distribution to rCoB (*77*, *78*). The results for the global ocean model are in the main text. By splitting the collection of metagenomes according to their Longhurst biome (*24*), we obtained 270 westerlies metagenomes, 225 trades metagenomes, 52 coastal metagenomes and 42 polar metagenomes. A distinct picture emerged by re-evaluating drivers of rCoB at a regional scale.

Marine biomes (except for the coastal biome) follow an absolute latitudinal gradient, ranging from both poles to the equator (Figure 1-B). Hence, we expected a decrease in the contribution of temperature when stratifying samples by biome. Biome-specific models revealed that: (i) Coastal biome is mainly driven by nutrients lead by phosphate. (ii) Westerlies biome communities displayed a balance between temperature and nutrients influence mainly driven by phosphate (the only nutrient with a p-value $<0.05$). (iii) Trades communities follow the same trends as Coastal, mainly influenced by nutrients even though silicate is the only significant nutrient of the model. (iv) No variables were significant for Polar biomes, maybe due to the low number of data, even if a clear correlation exists between temperature and rCoB (Figure S5). Altogether, these results at regional scale reveals that nutrients effects predominate where temperature tends to be stable across seasons(*26*).

<u>Supplementary figures:</u>

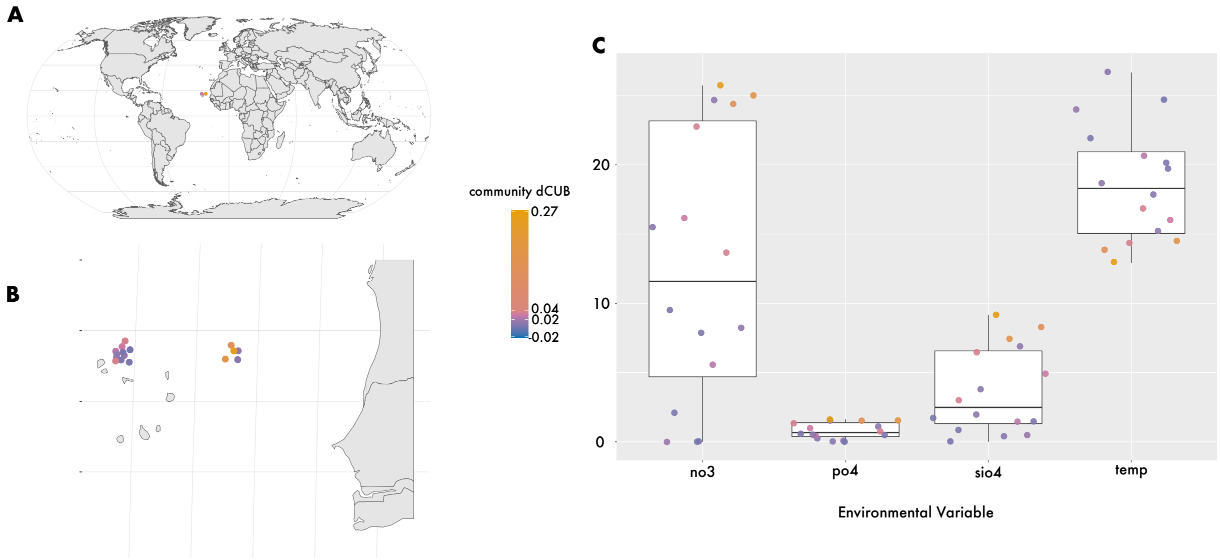


*Figure S1 :* **Cabo Verde high rCoB is link to local high nutrients concentration and low temperatures, another example of the low temperature high nutrient correlation.**

**A.B.** Localisation of the Cabo Verde communities plotted in this panel. Note that a small jitter was added on B to distinguish communities with similar coordinates but different depths*.* **C.** Environmental values of communities in the vicinity of Cabo Verde islands. temp = temperature. po4, no3 and sio4 = phosphate, nitrate and silicate concentrations. Note that metagenome with the maximum rCoB on these plots appears to be strongly under the influence of one prokaryote.  Alone it accounts for 65% of the total rCoB. This prokaryote is labelled MGYG000453028 and belongs to the *Bermanella* genus.

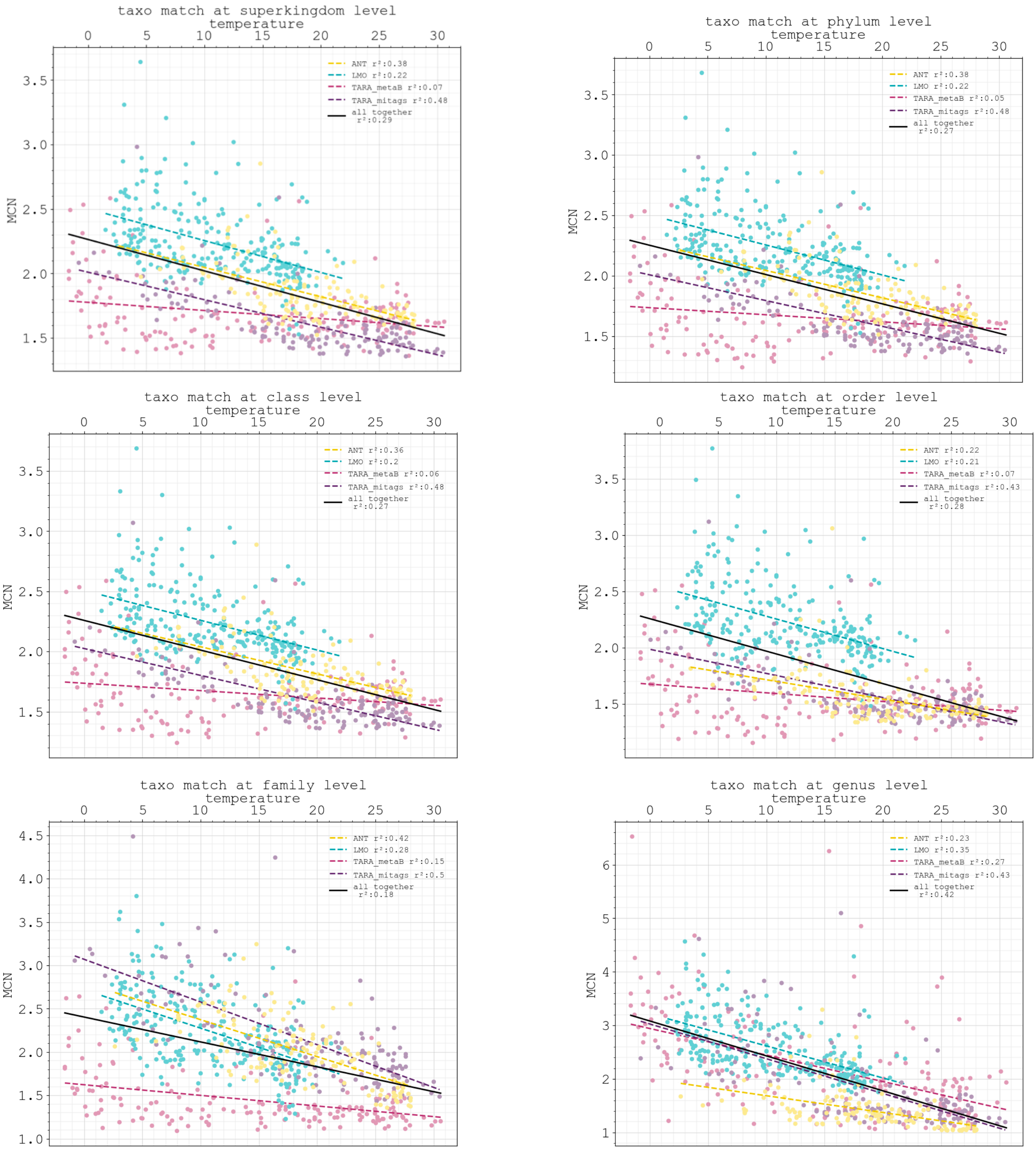


*Figure S2:* **Matching MCN copy number at genus and below levels increases its correlation with temperature.**

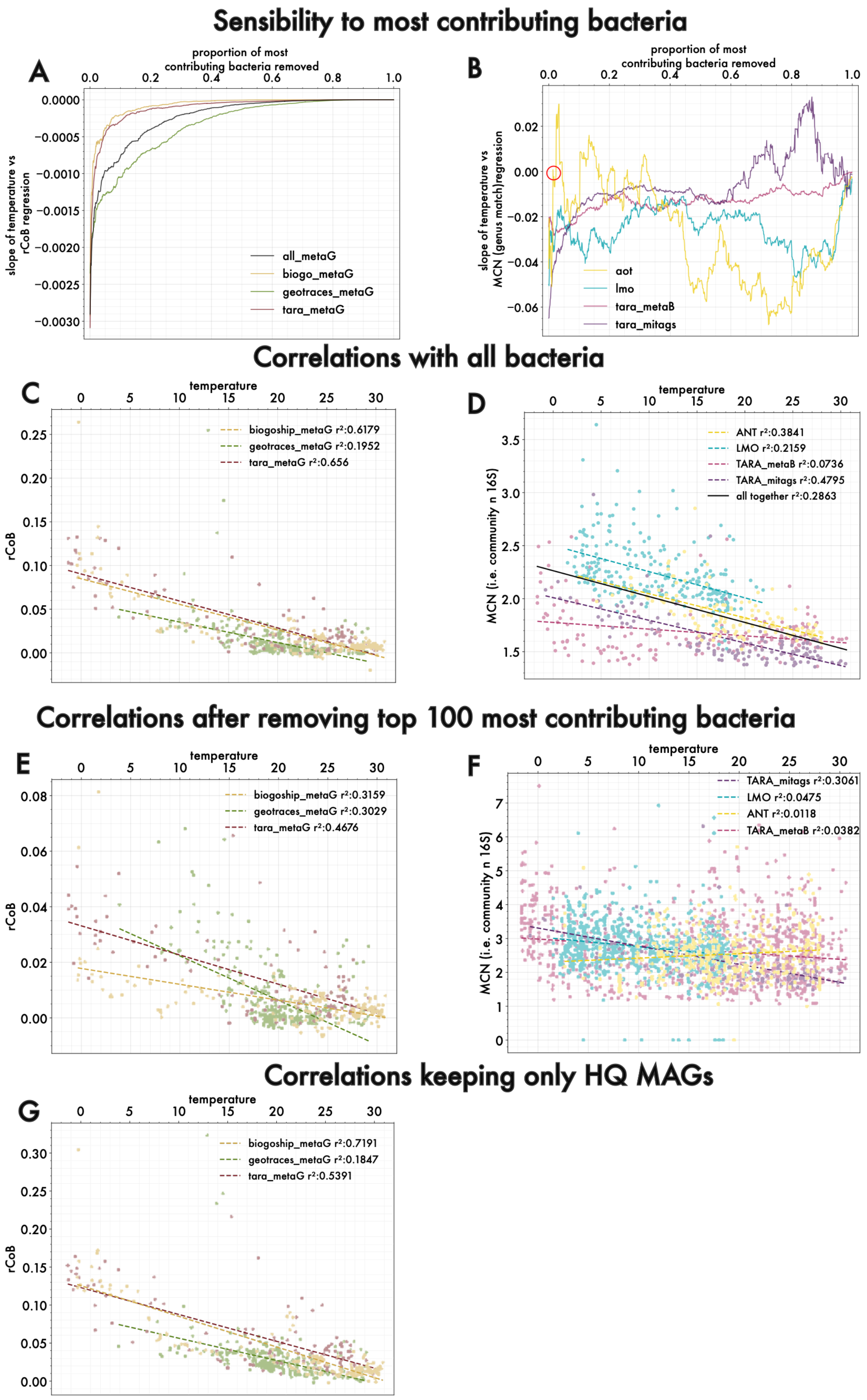
Sensibility to most contributing bacteria
A
proportion of most contributing bacteria removed
slope of temperature vs rCoB regression
all_metaG
biogo_metaG
geotraces_metaG
tara_metaG
B
proportion of most contributing bacteria removed
slope of temperature vs MCN (genus match) regression
aot
lmo
tara_metaB
tara_mitags
Correlations with all bacteria
C
temperature
rCoB
biogoship_metaG r²:0.6179
geotraces_metaG r²:0.1952
tara_metaG r²:0.656
D
temperature
MCN (i.e. community n 16S)
ANT r²:0.3841
LMO r²:0.2159
TARA_metaB r²:0.0736
TARA_mitags r²:0.4795
all together r²:0.2863
Correlations after removing top 100 most contributing bacteria
E
temperature
rCoB
biogoship_metaG r²:0.3159
geotraces_metaG r²:0.3029
tara_metaG r²:0.4676
F
temperature
MCN (i.e. community n 16S)
TARA_mitags r²:0.3061
LMO r²:0.0475
ANT r²:0.0118
TARA_metaB r²:0.0382
Correlations keeping only HQ MAGs
G
temperature
rCoB
biogoship_metaG r²:0.7191
geotraces_metaG r²:0.1847
tara_metaG r²:0.5391

*Figure S3:* **Comparison of rCoB and MCN properties.** *Sensibility to influent prokaryote removal of rCoB (A) and MCN (B). Correlation with temperature using all prokaryotes (C & D) in each community metric versus after removing the top 100 most contribution prokaryotes (E & F). Correlation keeping only High-Quality MAGs (contamination <5% completeness >90%) (G). Note the evolution of* $r^2$ *in the legends of all plots.*

When looking at the profiles of the contribution plots, it clearly appears that the slope between rCoB and temperature evolves in a much more predictable pattern than the slope between MCN and temperature (AB). For rCoB (A), when we progressively remove bacteria with high contribution, the slope progressively decreases toward zero, which is expected considering the fact that we remove first the bacteria tilting the slope. Moreover, the signal does not disappear after removing progressively the most contributing bacteria (D compared to C). For MCN, it is less predictable: it seems that omitting some bacteria can affect deeply the slope between MCN and temperature(B). For instance, in the ANT study, after removing only one hundredth of the most contributing bacteria of the study, the sign of the slope changes (B - yellow line, red circle). In a situation where the experiment would fail to detect a couple of bacteria, it would lead to opposite conclusions, suggesting an increase of MCN with temperature. The instability of MCN could be explained by the uncertainty taxonomic assignation of the copy number that do not reflect the reality, and by the fact copy number is discrete compared to the continuous property of rCUB.

Note that keeping strictly High-Quality MAGs do not modify the conclusions of temperature and rCoB correlation (G).

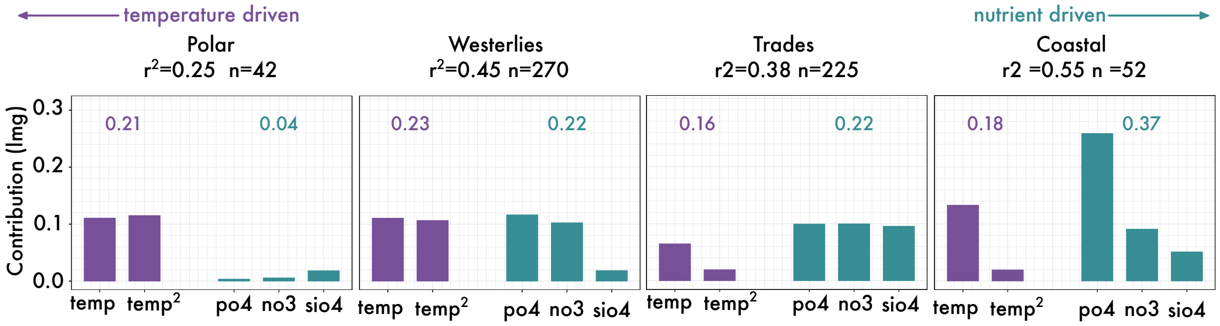


*Figure S4:* **Contribution analysis per Longhurst marine biome refines the understanding of temperature and nutrient concentration effect on explaining rCoB**. temp and temp$^2$ = linear and quadratic component of the polynomial fit between temperature and rCoB. PO4, NO3 and SiO4 = phosphate, nitrate and silicate concentrations.

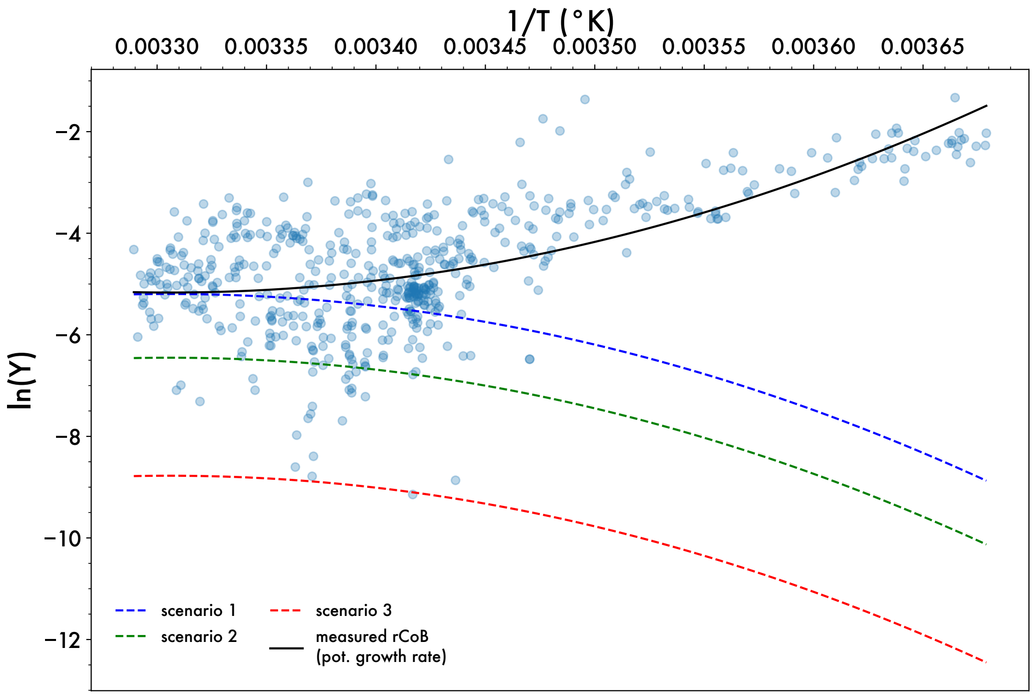


*Figure S5:* **Scenarios of differences between potential and effective growth rate in the ocean.** We plotted the log of our proxy of potential growth rate (pot. growth rate or rCoB) compared to the inverse of temperature (in Kelvin). We find a curve line as predicted by Arroyo et al. (*67).* Then, we plotted three scenarii for effective growth rate. Given the fact that effective growth rate cannot be higher than potential growth rate, and given the curvilinear response of effective growth rate to temperature, the three scenarios reach a maximum below the minimum of the effective growth rate. Whatever the scenario, the difference with potential growth rate is always higher on the right part of the plot, where we find cold communities, i.e., Polar communities. Hence, in these cold places, potential growth rate could be further from effective growth rate then it is in warm places. We could imagine that potential and effective growth rate match in warm communities, like plotted on scenario 1. Or we could imagine that effective never reaches potential growth rate, resulting in scenario 2 or 3 given on how big is the difference.

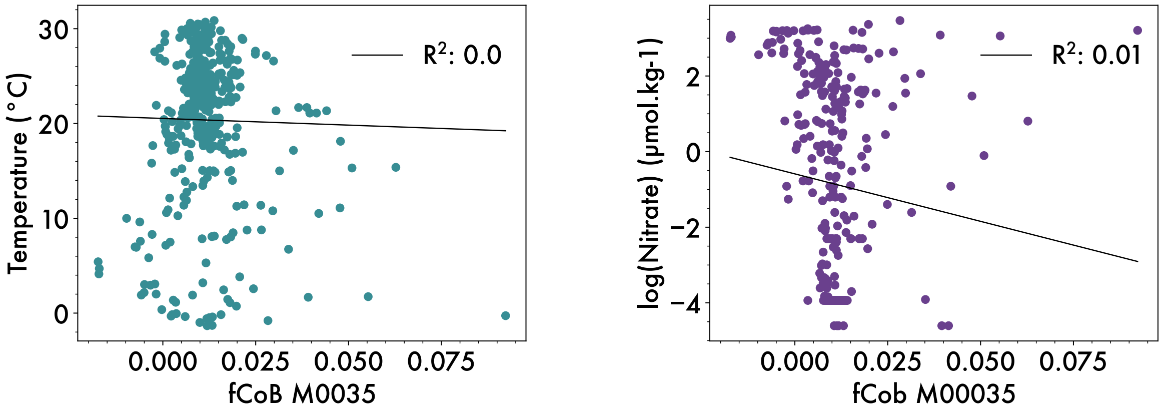


*Figure S6:* **M00035, a module that do not correlates with temperature nor** nitrate **concentrations**

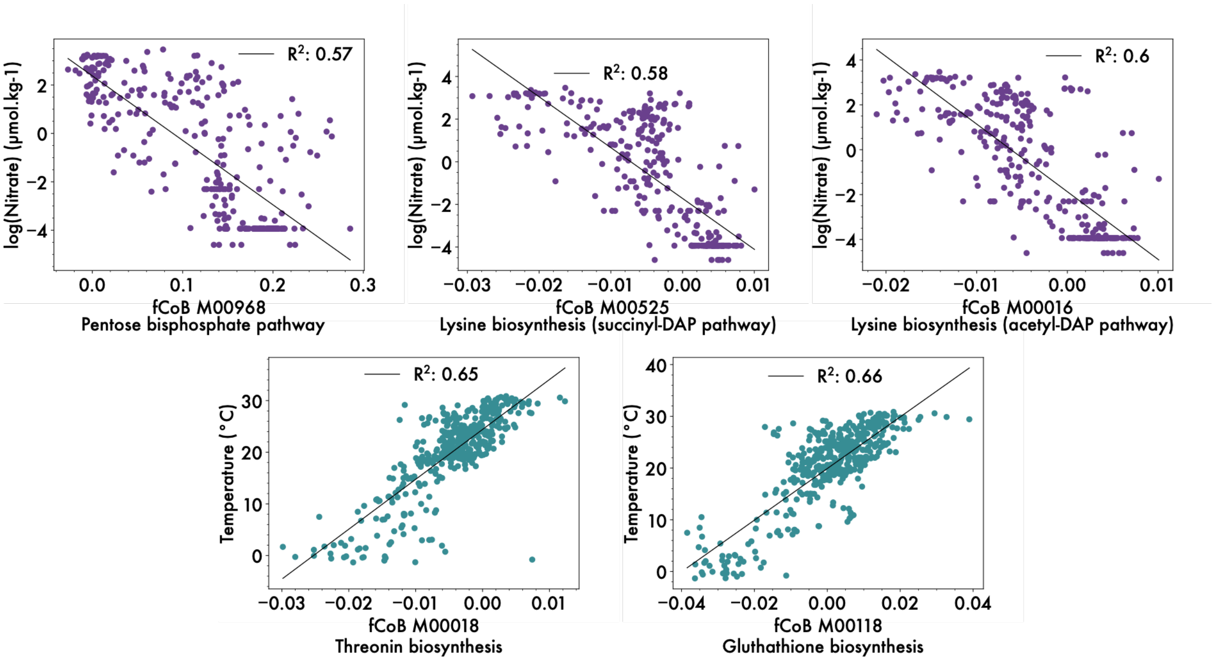


*Figure S7:* **Five fCoBs showing high global correlation with nitrate concentration or temperature.** They complement the two modules shown in the main text (M00136 and M00338)

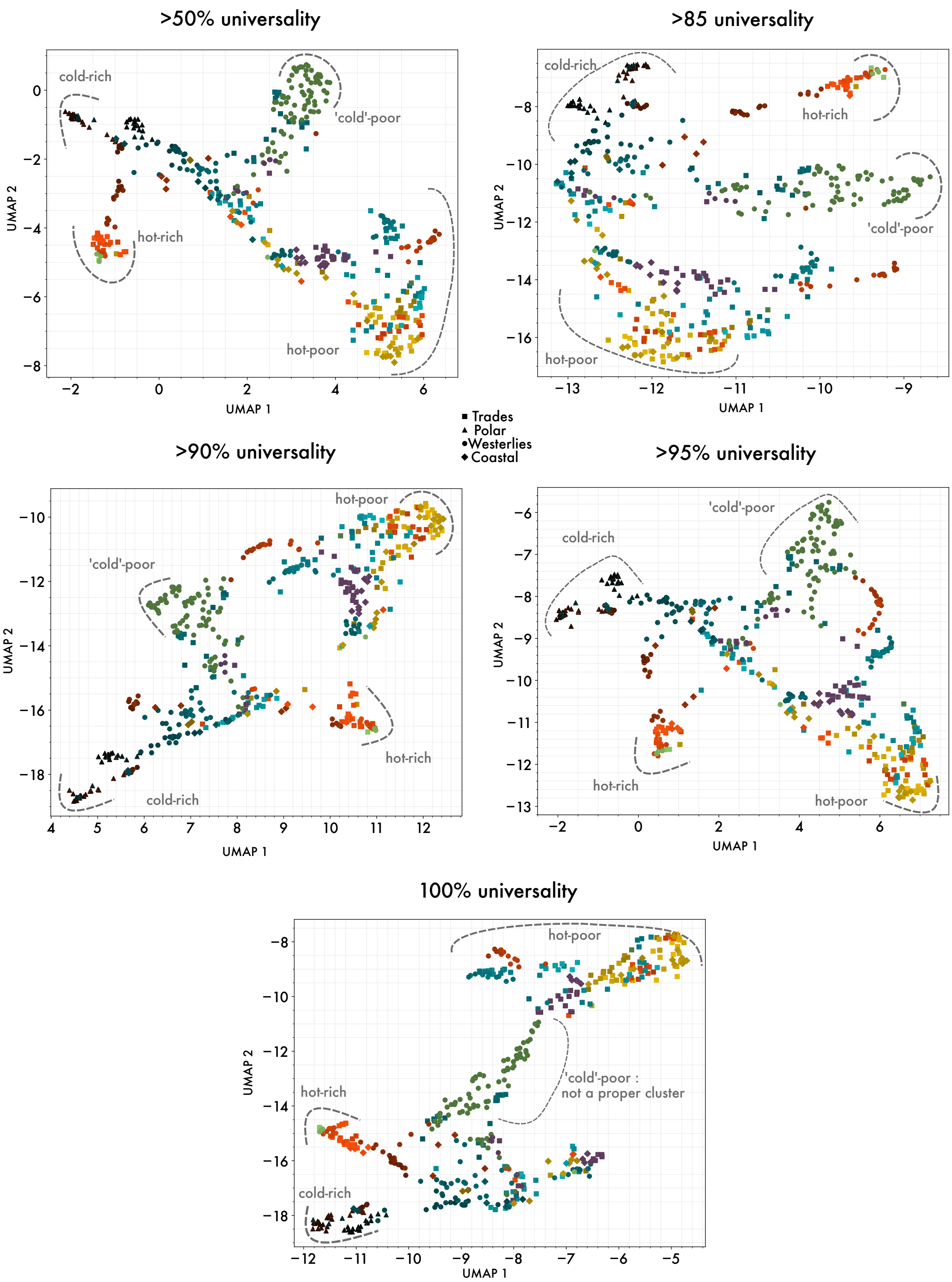


*Figure S8:* **UMAP projection with different thresholds on function universality.** We are always able to identify the four temperature-nutrient clusters until the threshold 95% on function universality, even if the limits of the clusters fade. At 100% the cold-poor cluster disappears, absorbed at the centre of the UMAP. This panel illustrates the trade-off on function universality: keeping only functions detected absolutely everywhere removes quasi-universal functions with relevant bias signature in answer to local environmental pressures.

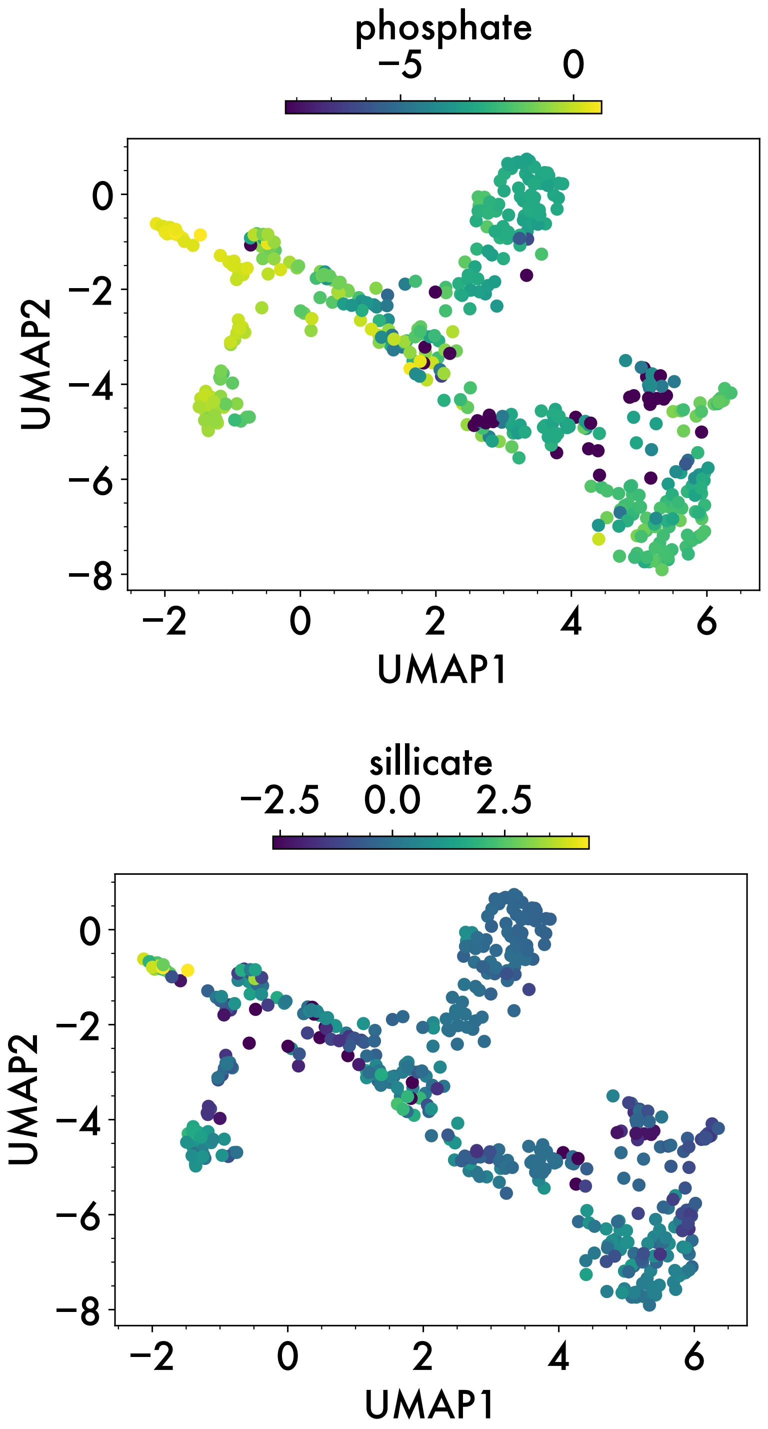


*Figure S9:* **Phosphate and silicate on UMAP follows nitrate concentrations.** Natural log scale was used, and when nutrient concentration was null, we imposed the colour to be equal to the log of the lowest non null value.

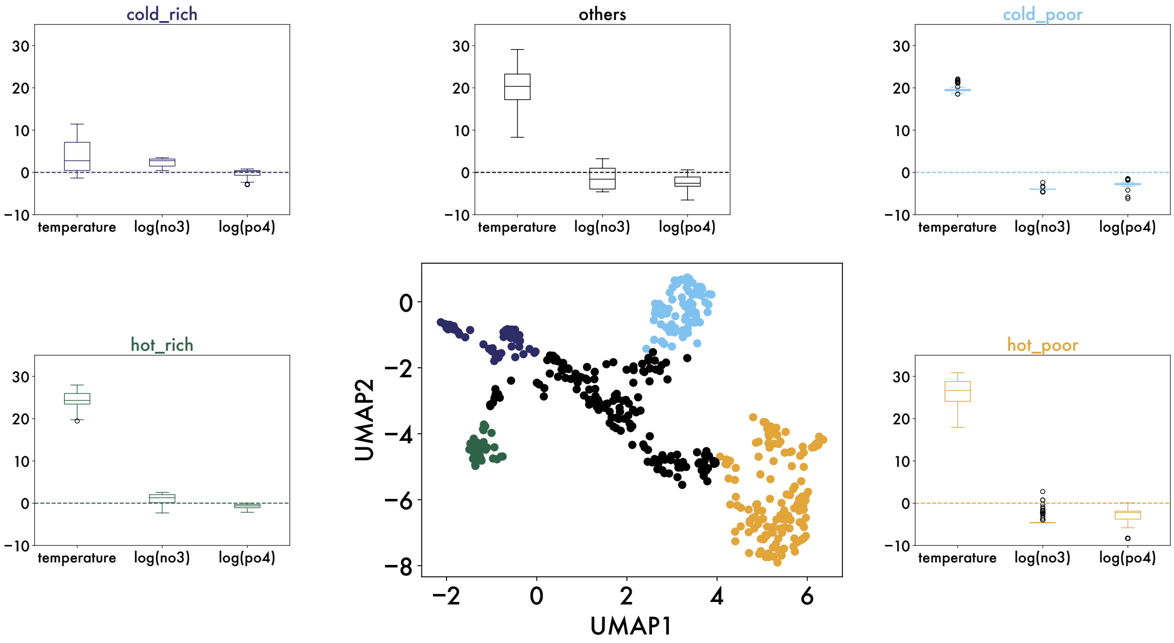


*Figure S10: **Clusters of the UMAP plot are structured by four distinct temperature-nutrient regimes.***

We grouped values of temperature and nutrient concentration per cluster (represented by different colours) to show that they all present a different combination of temperature and nutrient concentration values. To ensure visual comparability, the y scales are identical between the 4 boxplots.

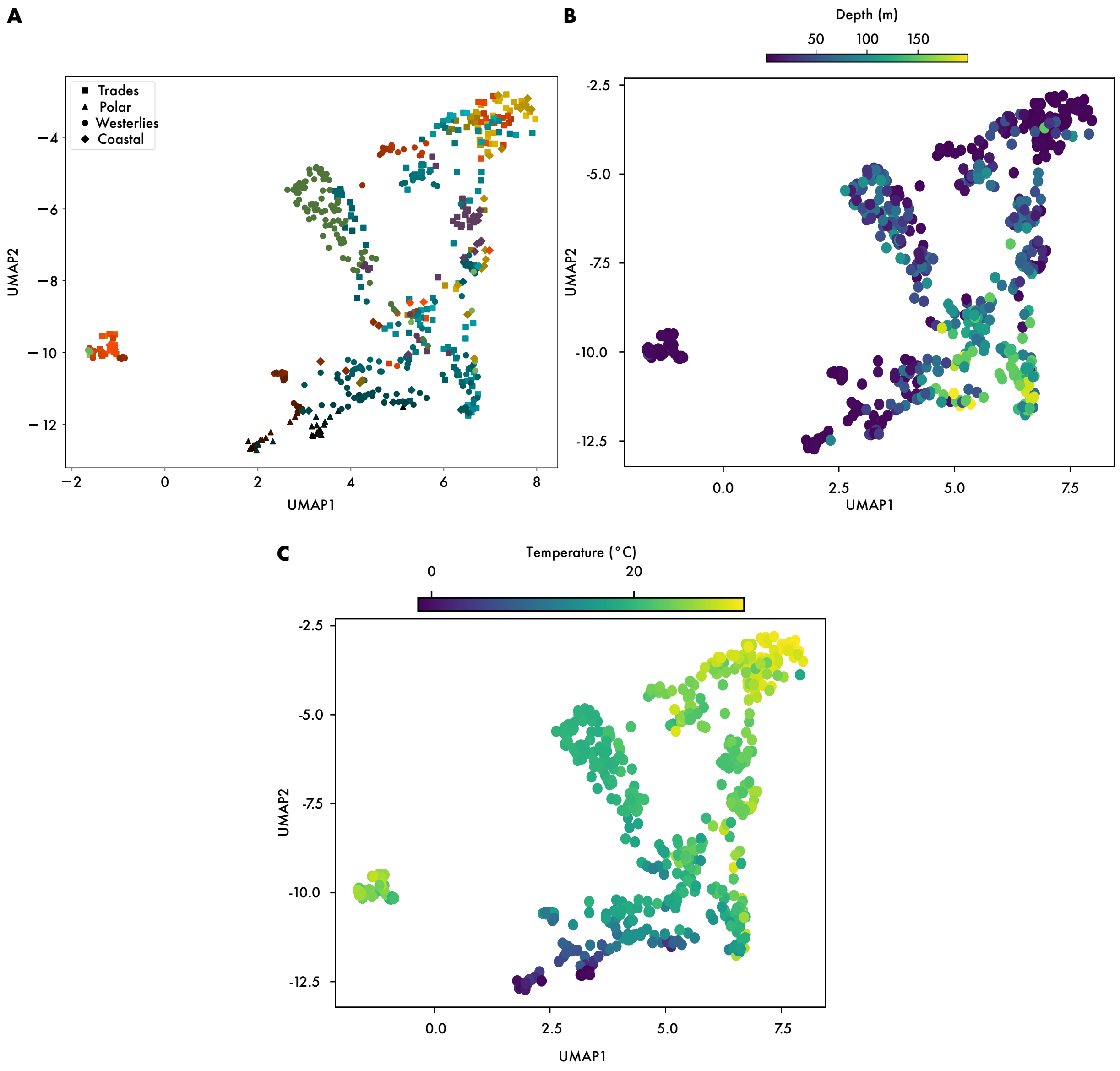


*Figure S11:* **Keeping deep (100-200m) communities modifies the UMAP by forming a depth-related cluster.**

**A**. UMAP plot when keeping all 589 metagenomes, including the deep 100-200m communities. **B.** The UMAP coloured by depth. **C.** The UMAP coloured by temperature

We can see a cluster formed on the right bottom part of the UMAP explained by depth (B). However, like seen on plot A, most of the deep metagenomes come from the Atlantic Ocean (from dark to vibrant blue). Hence, it complicates the understanding: is the cluster representative of the 100 to 200m depth layer, ocean wide? Or only from the Atlantic part? If the first hypothesis turns to be true, it would reveal a specific optimisation of functions influenced by depth directly or indirectly (e.g. light availability).

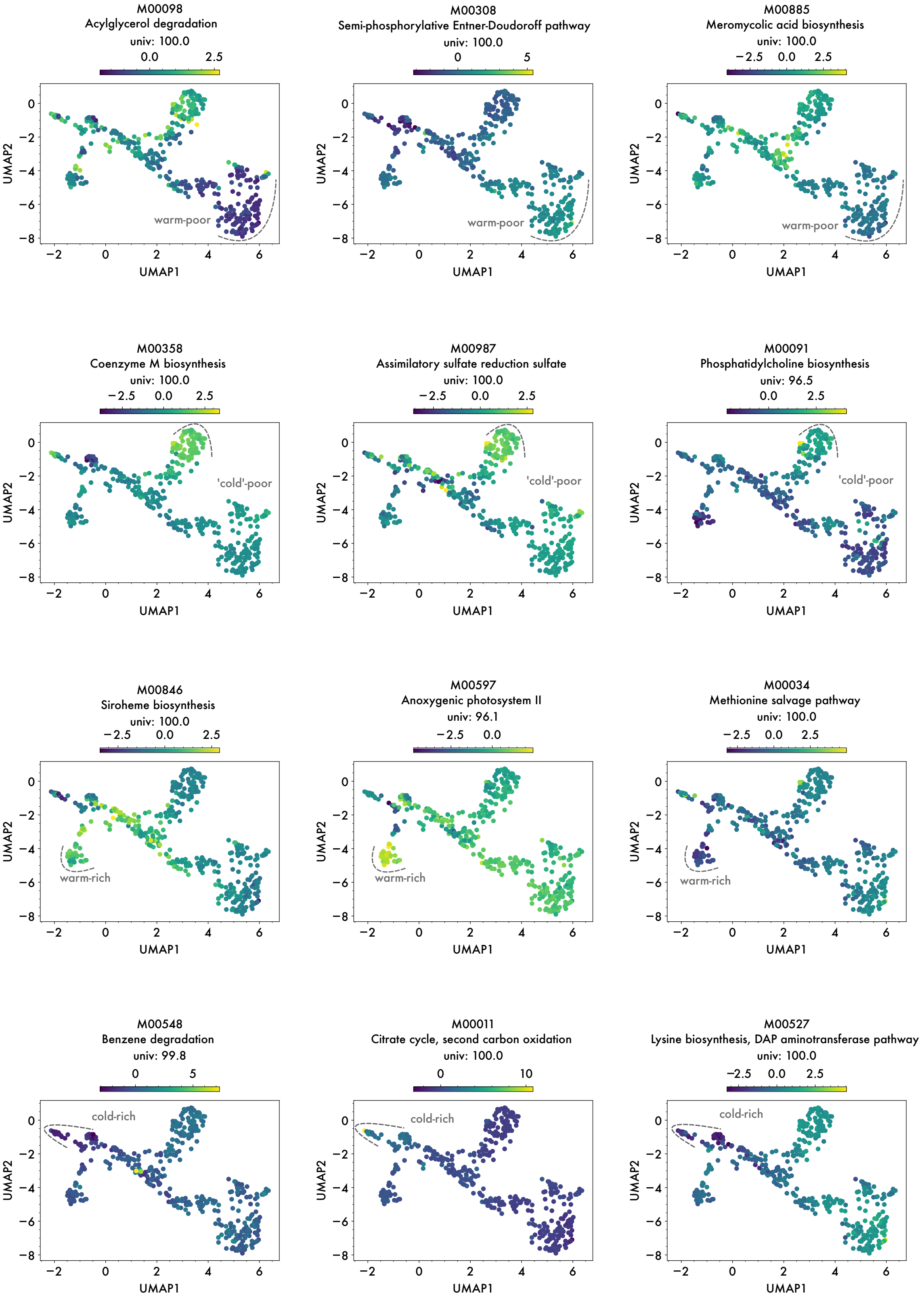


*Figure S12:* ***Visualizing typical functions driving each cluster on the UMAP****. One can visually see how each function shows unique values in the cluster it represents. Identified through gradient boosting, in order to separate the cluster from the 3 others. Among all the functions used to segregate clusters, we kept 3 visual representatives. Univ. =universality. It is the percentage of station of our study where the function was detected in.*

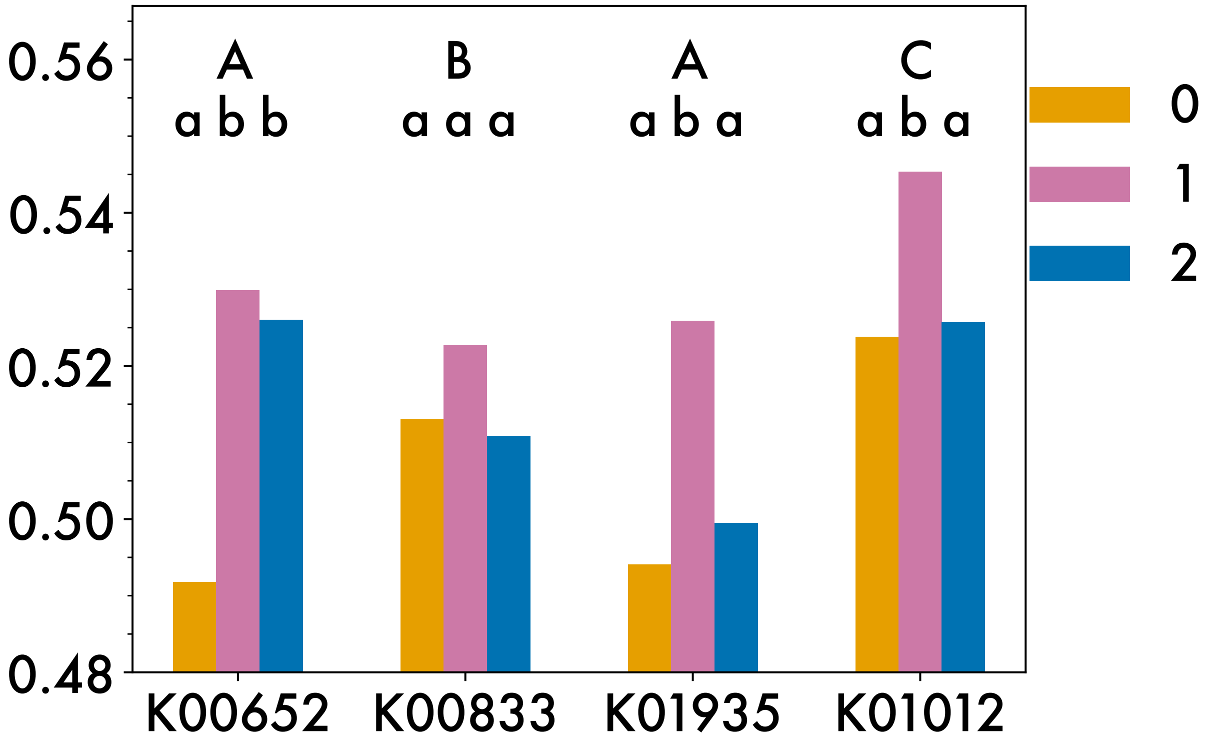


*Figure S13:* **Median bias of genes encoding the B7 biosynthesis pathway from 261 *Prochlorococcus* MAGs**.

Capital letters indicate significant differences in medians between genes of the pathway (all clusters taken together, permutation test, n=10000, p-value<0.05). Lower case letters indicate significant differences in medians between cluster, inside a given gene (permutation test, n=10000, p-value<0.05)
